# Super-Resolution Microscopy Reveals Structural Mechanisms Driving the Nanoarchitecture of a Viral Chromatin Tether

**DOI:** 10.1101/181016

**Authors:** Margaret J. Grant, Matthew S. Loftus, Aiola P. Stoja, Dean H. Kedes, Malcolm Mitchell Smith

## Abstract

By tethering their circular genomes (episomes) to host chromatin, DNA tumor viruses ensure retention and segregation of their genetic material during cell divisions. Despite functional genetic and crystallographic studies, there is little information addressing the three-dimensional structure of these tethers in cells, issues critical for understanding persistent infection by these viruses. Here, we have applied direct stochastic optical reconstruction microscopy (dSTORM) to establish the nanoarchitecture of tethers within cells latently infected with the oncogenic human pathogen, Kaposi's sarcoma-associated herpesvirus (KSHV). Each KSHV tether comprises a series of homodimers of the latency-associated nuclear antigen (LANA) that bind with their C-termini to the tandem array of episomal terminal repeats (TRs) and with their N-termini to host chromatin. Super-resolution imaging revealed that individual KSHV tethers possess similar overall dimensions and, in aggregate, fold to occupy the volume of a prolate ellipsoid. Using plasmids with increasing numbers of TRs, we found that tethers display polymer power-law scaling behavior with a scaling exponent characteristic of active chromatin. For plasmids containing a two-TR tether, we determined the size, separation, and relative orientation of two distinct clusters of bound LANA, each corresponding to a single TR. From these data, we have generated a three-dimensional model of the episomal half of the tether that integrates and extends previously established findings from epi-fluorescent, crystallographic, and epigenetic approaches. Our findings also validate the use of dSTORM in establishing novel structural insights into the physical basis of molecular connections linking host and pathogen genomes.

## Introduction

Kaposi’s sarcoma-associated herpesvirus (KSHV) is the etiological agent of Kaposi’s sarcoma, an inflammatory tumor of the skin and mucous membranes, as well as two B cell tumors, primary effusion lymphoma (PEL) and multicentric Castleman’s disease (1–3). KSHV infection is marked by long periods of latency when the virus expresses only a small number of essential genes (4). Among these is ORF73 that encodes LANA (5, 6), a protein of 1162 amino acids with multiple functions, including the tethering of viral episomes to host chromatin. The N-termini of LANA dimers bind to histones H2A/B and to chromatin-associated bromo-domain proteins (7–10). The C-termini of three LANA dimers bind to three adjacent 20-bp LANA binding sites (LBS 1, 2, and 3) located in each 801-bp TR (Fig. 1A) (11–15). There are an estimated 35 to 60 tandem TRs per viral episome (16, 17), arranged head-to-tail at a single location on the circular episome. The resulting tether, comprised of this cluster of TRs and bound LANA dimers, is essential for episomal maintenance within dividing cells and, presumably, persistence in patients (12, 18–21). Initial epifluorescence studies identified LANA as nuclear punctae recognized by serum antibodies from KSHV infected patients (22–25). Ten years later, Adang et al. used a combination of flow cytometry and qPCR to demonstrate a direct proportionality between the number of LANA punctae and the amount of viral DNA, leading to the conclusion that each nuclear dot represented a single viral episome (26). Previous studies examining these punctae showed association of LANA with mitotic chromosomes at the resolution of epifluorescence microscopy (12, 27). Kelley-Clarke et al. later suggested preferential localization of these LANA punctae to centromeric and telomeric regions on metaphase chromosomes (28). Such studies have provided a solid foundation for further inquiry into the nature of this tethering mechanism.

**Fig. 1.**
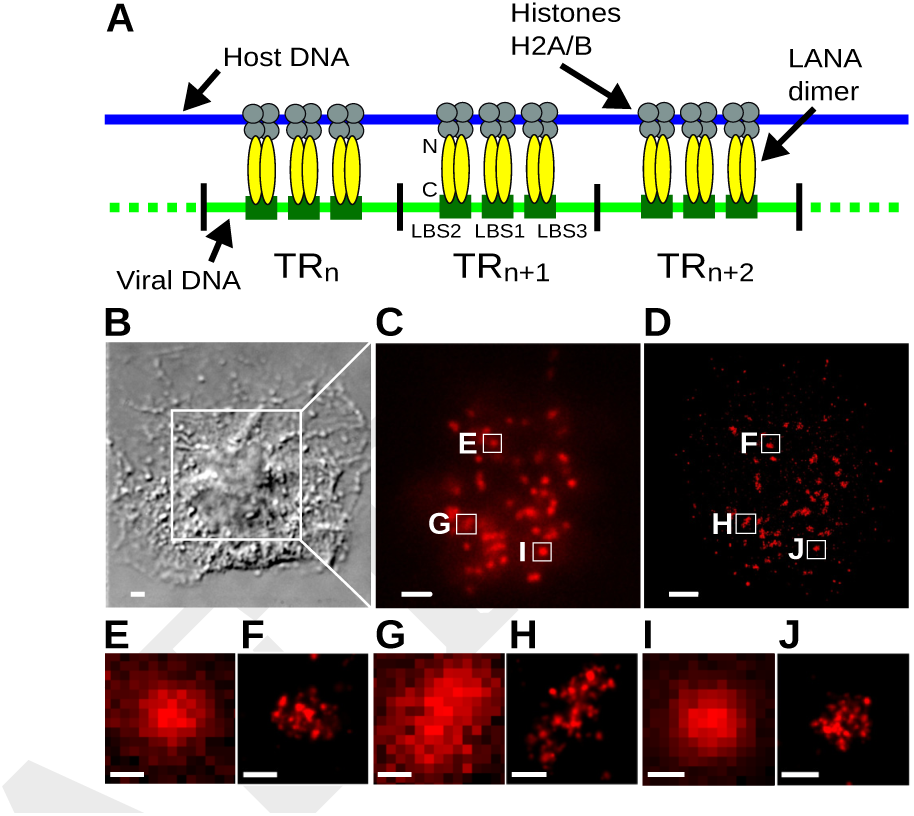
dSTORM offers improved image resolution of LANA tethers. (*A*) Schematic depiction of the anatomy of a KSHV LANA tether. LANA dimers (shown in yellow) bind via their N termini to histones H2A/B (grey). The C termini bind with sequence specificity to three LANA binding sites (LBSs) located on each terminal repeat (TR). (*B*) An anti-LANA stained BCBL-1 cell imaged by differential interference contrast microscopy. (*C*) The boxed region in (B) viewed by conventional epifluorescence microscopy. (*D*) The same field of view in (*C*) imaged by dSTORM microscopy. (Scale bars for B-D = 2 *μ*m), (*E*, *G*, and *I*) The three LANA tethers boxed in (C) are enlarged for better visualization. (*F*, *H*, and *J*) The three LANA tethers boxed in (*D*) are enlarged for comparison with their paired standard epifluorescence images (*E*, *G*, and *l*). (Scale bars for *E*-*J* = 250 nm).

Many fundamental features of full-length tethers in cells have remained elusive due to the resolution constraints of epifluorescence microscopy. While X-ray crystallography has resolved the structures of an N-terminal 23 amino acid LANA peptide bound to nucleosomes, and a C-terminal 139 amino acid peptide in complex with LBS1 (7, 15), questions remain as to the architectural features of a full-length tether. It is unknown how TR chromatin folds and whether this is consistent among tethers, both within and across cell lines. The close packing of nucleosomes and presence of heterochromatin protein 1 (HP1) at TRs found by others implies that the region is characterized as heterochromatin, whereas the presence of histone acetylation found by several groups suggests active chromatin (29–32). Questions also remain as to how LANA is distributed across the TRs, whether it attains full occupancy on all three LBSs, and whether supernumerary LANA dimers form large complexes at each TR to entrap episomal DNA loops (33). There is also no information indicating whether the number of TRs bound by LANA reaches a plateau. LANA dimers might bind to a maximum number of TRs in a plasmid or episome, regardless of the number of total TRs available. Finally, it is not clear how all of these elements influence tether folding and the ability to effectively bind to both viral and host genomes. To address these questions, we applied super-resolution microscopy to study KSHV infected cells with the goal of generating architectural data on full-length KSHV tethers.

## Results

### The Nanoscale Dimensions of LANA Tethers are Uniform and Consistent Across Different Cell Types

In principle, since LANA binds both host nucleosomes and episome DNA, the packaging of TR chromatin could be determined primarily by the state of the host chromatin, by features intrinsic to the viral episome, or both. To distinguish among these alternatives, we examined the tether morphology of the same viral isolate in two different cell types. We found that dSTORM resolved each epifluorescent LANA dot into a cluster of individual emissions from fluorophore-labeled anti-LANA mAbs (Fig. 1B-J). We interpret these emissions as blinks within a Gaussian localization volume from each conjugated dye molecule present at approximately three dye molecules per antibody (Materials and Methods), up to six antibodies bound per LANA dimer (see discussion of coiled-coil domain, below), and up to three LANA dimers per TR. Since all the episomes from the same virus have the same number of TRs (17, 34), and each TR is 801 bp, we asked whether the polymer architectures revealed by dSTORM would be similarly consistent across multiple tethers. We examined 65 individual LANA tethers from BCBL-1 cells (a KSHV-positive PEL line) and 34 from SLKp/Caki-1p cells, a human epithelial line exogenously infected with KSHV derived from BCBL-1 (35, 36). We found that while each tether presented a unique shape, their overall dimensions were similar in the two cell lines (Fig. 2A and B). To quantify this similarity, we aligned clusters along their three principal axes and calculated their dimensions and radii of gyration, *R_g_* (Materials and Methods), shown in Fig. 2C and D. The median *R_g_* for tethers in BCBL1 cells was 341 nm, 95% CI [309-353] and in SLKp cells it was 368 nm, 95% CI [323, 405]. There was no statistical difference between the *R_g_* of BCBL-1 and SLKp tethers (*P* = 0.26, Mood’s Median Test). The similarity of these parameters in a single viral strain in two different cell types shows that the polymer folding of KSHV LANA tethers is likely to be intrinsic to the viral episome itself, rather than being a result of the cellular milieu.

**Fig. 2.**
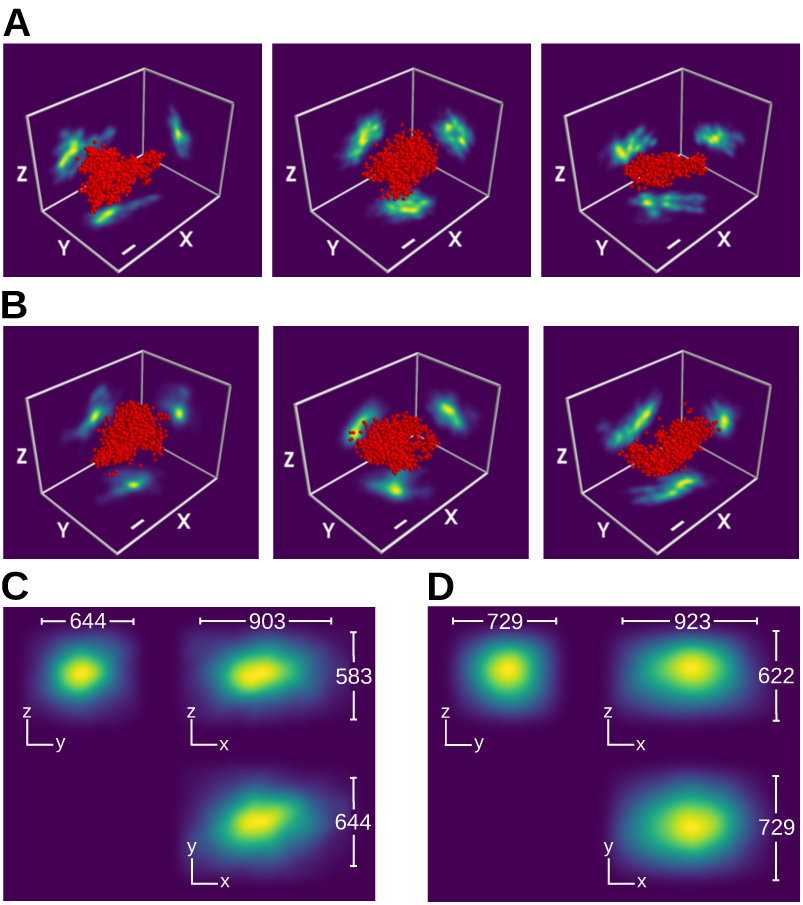
KSHV LANA tethers have consistent dimensions within and between cell types. (*A*) Three-dimensional projections of three representative LANA tethers from asynchronous BCBL-1 cells. Red spheres depict the Cartesian coordinates of individual photon emissions, with their radii scaled to the median axial localization precision of 30 nm. (*B*) Three representative LANA tethers from asynchronous SLKp cells, shown as in (*A*). (Scale bars for A and B = 250 nm.) (*C*) LANA tethers from BCBL-1 cells (N=65) were oriented by principal axes and aligned at their centroids. The compilation is rendered as Gaussian emissions with 30 nm localization precision and presented from three architectural viewpoints, indicated by the X, Y, and Z directional axes. Median dimensions are shown by the bars. (*D*) LANA tethers from SLKp cells (N=34) superimposed and rendered as in (*C*).

### LANA Occupancy Scales with the Number of TRs

We postulated that if the tethers visualized by dSTORM reflect the underlying TR architecture, then their structural parameters should change systematically with different numbers of TR elements. To challenge that hypothesis, we examined synthetic tethers with different numbers of TRs. We co-transfected BJAB cells, a KSHV-negative human B cell line, with two plasmids, one encoding LANA and the other containing 2, 5, 7, or 8 tandem TRs arranged head-to-tail as in the native episome (Fig. S1A and B). In addition, we infected BJAB cells with BAC16 virus that we determined contains approximately 21 TRs (Fig. S1C and D). Transfection with a LANA-encoding plasmid and a plasmid lacking any functional TR sequence (p0TR) resulted only in diffuse nuclear staining by epifluorescence (Fig. S2A), consistent with earlier work (12, 37). In contrast, individual tethers in cells harboring plasmids with 2, 8, or 21 TRs showed an increase size with increasing number of TRs (Fig. 3A-C). This increase was accompanied by a corresponding linear increase in the number of dSTORM fluorophore emissions and, therefore, anti-LANA Ab binding (Fig. 3D). Our observation that an increase in TR number results in a proportional increase in LANA binding argues against a plateau for TR occupancy per plasmid or episome, at least over the range of 1-21 TRs.

**Fig. 3.**
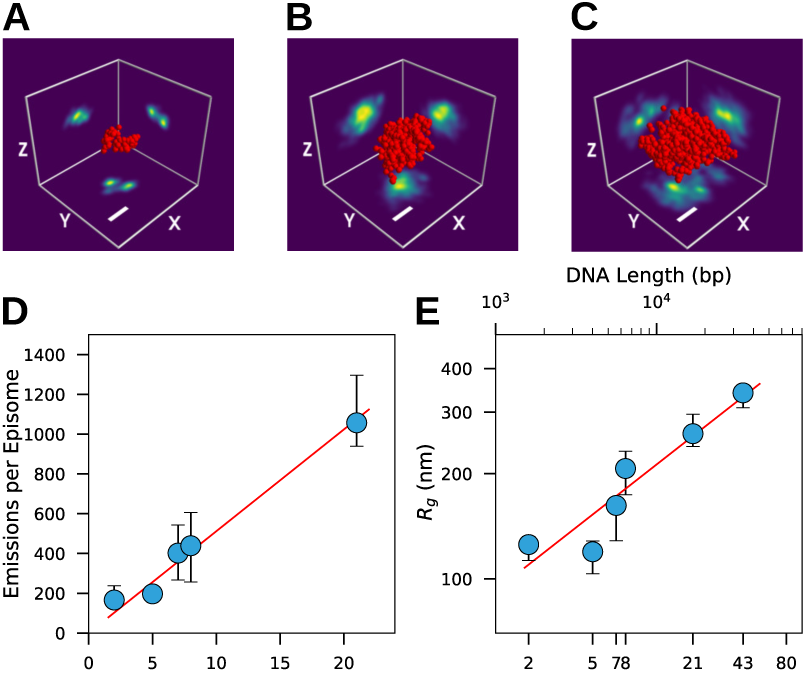
LANA tethers show polymer-like scaling as a function of the number of TRs. (*A-C*) The 3D-projections of representative LANA tethers from a cell transfected with p2TR (*A*), p8TR (*B*), or BAC16 with 21 TRs (*C*) are shown at identical scales as described in Fig. 2. (Scale bar for *A-C* = 250 nm). (*D*) The linear relationship between the median number of emissions and the number of TRs per tether is shown for p2TR (N=28), p5TR (N=39), p7TR (N=13), p8TR (N=17), and BAC16 (N=32) (*R^2^ =* 0.99). (*E*) The log of median *R_g_* data for the tethers in (*D*) and BCBL-1 KSHV episomes (43 TRs, N=65) are shown as a function of the log of the tether DNA lengths (top axis), or the log of the TR number (bottom axis). The regression (solid red line) through tethers with known numbers of TRs (blue circles) was used to calculate the scaling exponent *c* = 0.36 ± 0.07 (*R*^2^ = 0.94). Error bars represent 95% cls.

### Tether Polymer Conformation has the Characteristics of Active Chromatin

At present, there is conflicting evidence regarding the conformational state of TR chromatin, which exhibits features of both repressed and active conformations. On one hand, TR nucleosomes are tightly packed in a typically “closed” conformation (31). Furthermore, LANA recruits the histone H3 methyltransferase SUV39H1 and establishes heterochromatin protein 1 (HP1) binding to the tether chromatin (30). On the other hand, TR chromatin is also modified by histone hyperacetylation characteristic of active chromatin (29, 31, 32), and the tether interacts with bromodomain protein BRD4 (9, 38).

To address the conformational state directly, we took advantage of the fact that the radius of gyration, *R*_g_, of a chromatin polymer follows the power-law relationship, *R_g_ ∝ L^c^*, where *L* is the DNA length in bp and *c* is the scaling exponent. This scaling parameter, *c*, is a sensitive measure of canonical active, inactive, and repressed states of mammalian chromatin (39). By plotting *R_g_* against the tether DNA length in bp we determined the power-law scaling exponent for the LANA-bound TR chromatin to be *c* = 0.36 ± 0.07 (Fig. 3E). This exponent approximated the value of 0.37 ± 0.02 found by Boettiger and co-workers for active chromatin and is distinct from values for inactive (*c* = 0.30 ± 0.02) or repressed (*c* = 0.22 ± 0.02) chromatin (39). We obtained similar results for TR plasmid scaling in COS-7 cells (Fig. S3A). This finding clarifies the long-standing paradox surrounding the chromatin folding state of the TR region.

### Adjacent TRs are Arranged with a Specific Asymmetric Architecture

Closer analysis of the dSTORM emissions from individual p2TR tethers revealed that they frequently gave rise to two distinct, resolvable clusters (Fig. 3A) and alignment of the complete set of 28 p2TR data (Materials and Methods) reinforced this separation (Fig. 4A). Individual clusters were characterized by an *R_g_* of 90 nm, 95% CI [82, 96], which is similar to the *R_g_* of 86 nm predicted by extrapolation of the power-law function to a single TR (Fig. 3E). Thus, we interpreted the two clusters as the signals emanating from two adjacent TRs. We obtained similar results for p2TR dSTORM images acquired in COS-7 cells (Fig. S3B). The p2TR tethers, individually or as an aligned composite, showed striking asymmetry. They comprised flat pairs of prominently elongated ellipsoids whose centroids were spaced apart by 174 nm, 95% CI [152, 185] (Fig. 4B). Moreover, the long axes of the two clusters intersected to form a 46° angle (Fig. 4B), demonstrating a specific differential placement of the individual TRs within the p2TR structure. These parameters provided strong constraints on the underlying folding of the p2TR chromatin tether.

**Fig 4:**
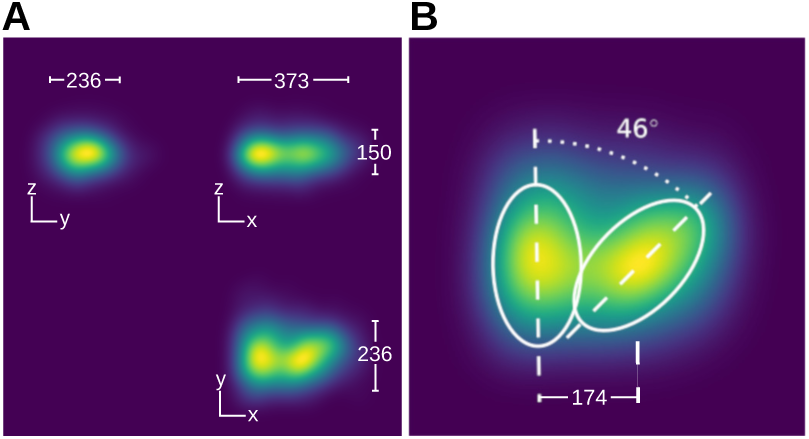
p2TR together show two distinct clusters LANA Ab that from a46° angle between their long axes. (*A*) Compilation of emission data from 28 p2TR tethers; emissions are rendered as in Fig. 2. Median dimensions (nm) of the combined data are shown. (*B*) Approximately 2X magnified view of the Y-X projection from A demonstrating the 46° angle formed between the two clusters.

### 2TR Images Support the Prediction of a LANA Coiled-Coil Domain

To begin to interpret the 2TR tether images at the molecular level, we evaluated the structure of LANA dimers N-terminal to the LBS binding domains, for which previous X-ray crystal structures are known (10, 15, 38). Examination of the primary protein sequence of LANA with `Paircoil2` (40) revealed a segment strongly predicted to form a coiled-coil (Fig. 6A; *P* < 0.025).

To test for the presence of the coiled-coil, we measured the number of single molecule emissions detected by dSTORM for LANA tethers stained with LN53-A647, which recognizes the tetrapeptide epitope EQEQ (41). There are 22 such epitopes within the LANA protein sequence and this number of antibodies would yield over 18,000 dSTORM emissions per LANA tether (Materials and Methods). However, 19 of the most C-terminal of these epitopes would be embedded within the putative coiled-coil region of LANA, occluding antibody recognition. The remaining three epitopes just N-terminal to the coiled-coil would yield on the order of only 2,500 emissions per tether. Indeed, we routinely recovered fewer than 2,000 emissions per tether, consistent with only 2 to 3 functional epitopes per LANA protein, providing strong empiric data for the predicted coiled-coil. Furthermore, the positioning of these available epitopes at the N-terminal end of a rigid coiled-coil domain is supported by the separation between the two emissions clusters in the 2TR images. A molecular model of LANA dimers incorporating these features is illustrated in Fig. 6.

### LANA-Imposed DNA Bending and Nucleosome Translational Positioning Are Predicted to Direct TR Chromatin Folding

Although the DNA of each TR is known to be bent by LANA dimer binding (15, 42, 43), and to be occupied by four nucleosomes (31), the constraints these factors impose on tether architecture are currently unknown. To address this question, we varied the parameters of DNA bending and nucleosome positioning in a set of molecular models (see Fig. 7).

**Fig. 7.**
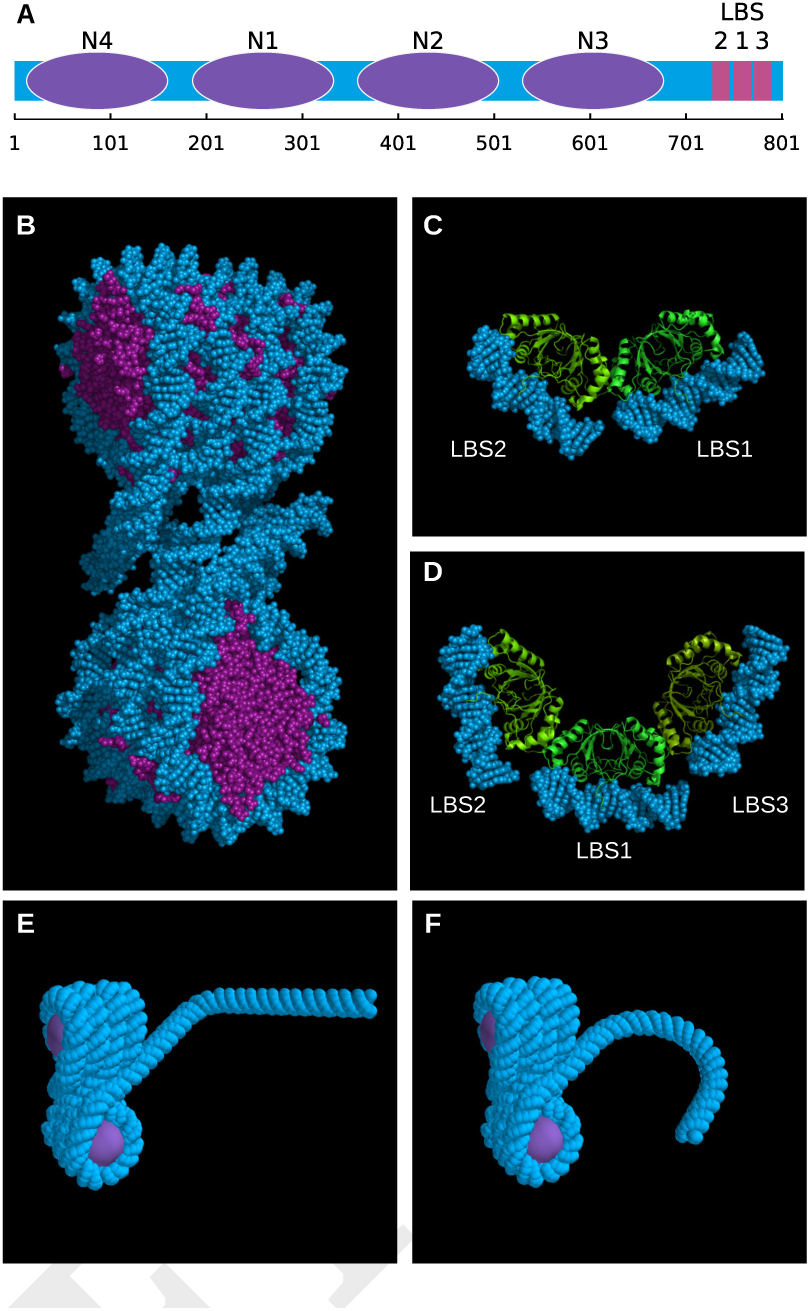
Modeling LANA dimers. (*A*) Schematic diagram of a single 801-bp TR showing localization of four nucleosomes (purple ovals) and three LBSs (magenta rectangles) on the DNA (blue). (*B*) Structure of DNA (blue) wrapping around four nucleosomes (purple), adapted from the tetrasome crystal structure (PDB 1ZBB). (*C*) Structure of two LANA dimers (green) binding to two adjacent LBSs (blue) of a single TR, showing the 110 degree DNA bend formed by protein binding. (*D*) Structure of three LANA dimers bound to all three LBSs in a TR, demonstrating the nearly 180 degree bend in the bound DNA. (*E*) Model of the DNA (blue) in a single TR, including the wrapping of four nucleosomes (purple), showing the 110 degree bend formed when only two adjacent LBSs are bound by LANA. (*F*) Model as in *E* but showing the 180 degree bend resulting from all three LBSs bound by LANA.

We modeled the 113 bp of nucleosome-free DNA that contains the three LBS sites in two modes. In the first, both LBS1 and LBS2 are occupied by LANA dimers, while LBS3 is unoccupied, inducing a 110° bend in LBS DNA (Fig. 7C), as previously determined by Wong and Wilson (42). In the second mode, LBS1, LBS2, and LBS3 are all occupied by LANA dimers, inducing a bend of approximately 180° (Fig. 7B), as predicted by Hellert et al. (15). For the remaining 688 bp of TR DNA occupied by four nucleosomes, we adapted the crystal structure of a synthetic tetrasome (PDB 1ZBB) (44) because of its similarity to the spacing of the nucleosomes mapped to the TR (31) (Fig. 7B). We digitally joined the tetrasome DNA to the LBS DNA, bent at either 110° (Fig. 7E) or 180° (Fig. 7F). We varied the position of the joint by a single base pair at a time over a 9 bp span, designated phases 1-9, to model the the effect of nucleosome translational positioning on tether architecture through an entire turn of a DNA helix (Fig. 8A and E).

**Fig. 8.**
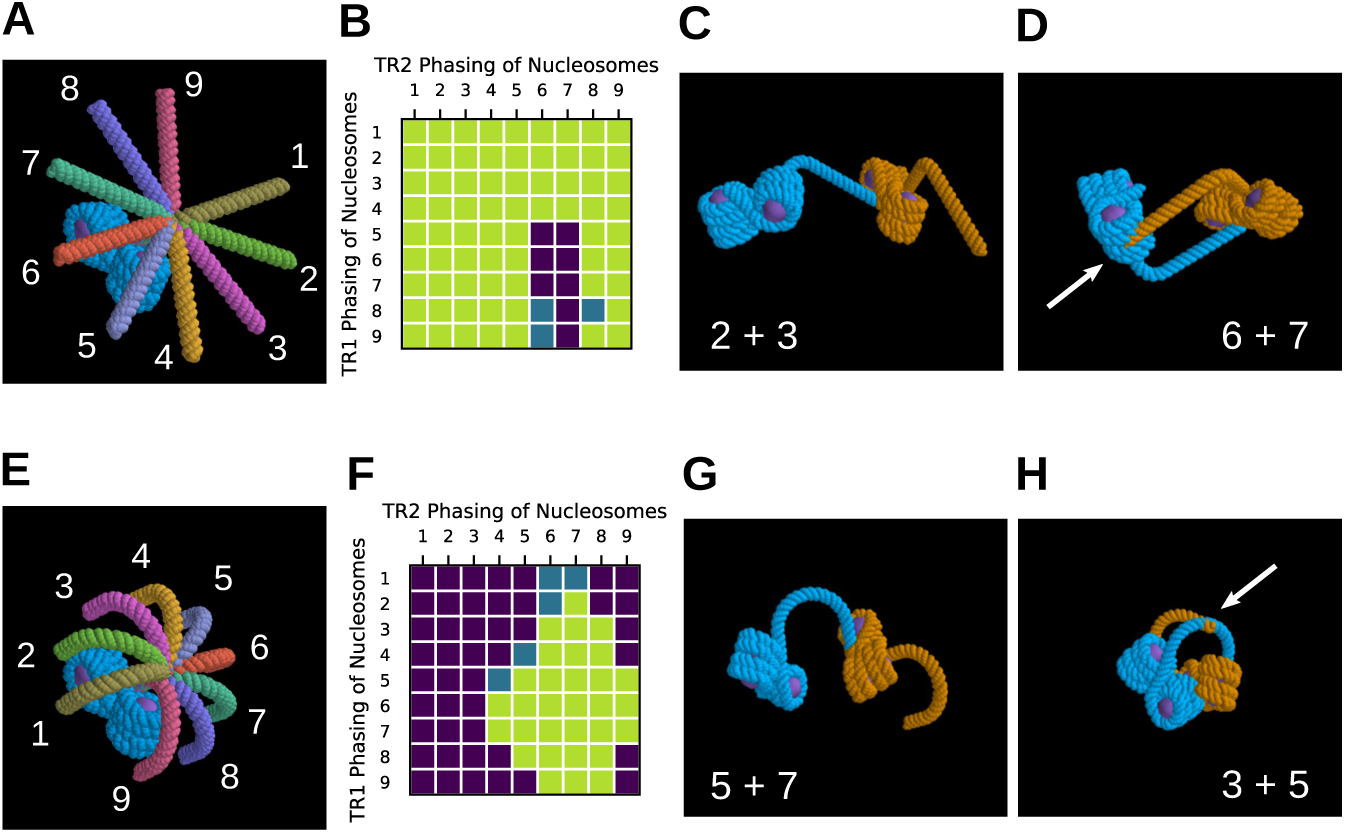
Modeling 2TRs. (*A*) A DNA-wrapped tetrasome (blue) displaying the rotational effects of the nine theoretically possible nucleosome phases on the position of the LBS-containing DNA fragment emerging from a single TR in which only two LBSs are bound by LANA (numbered 1 through 9). (*B*) A chart depicting the combinations of nucleosome phasing for two sequential TRs (TR1 and TR2), in which only two LBSs are bound by LANA. The nucleosome phasing results in “disallowed” arrangements (purple squares), wherein the elements face steric hindrance, “permitted” arrangements (green squares), in which there is no steric hindrance, and arrangements with likely interference (blue squares). (*C-D*) An example of a “permitted” phasing combination from B (TR1 = phase 2, TR2 = phase 3) is shown in *C*, in contrast to a “disallowed” phase (TR1 = phase 6, TR2 = phase 7) presented in *D*. Regions of steric hindrance are marked with white arrows. (*E*) The nine theoretically possible phases of the LBS-containing DNA fragment emerging from a TR in which all three LBSs are bound by LANA. (*F*) A chart depicting nucleosome phasing combinations for two sequential TRs in which all three LBSs are bound by LANA. Colors are as in B. (*G-H*) An example of a “permitted” phasing combination from *F* (TR1 = phase 5, TR2 = phase 7) is shown in *G*, in contrast to a “disallowed” phase (TR1 = phase 3, TR2 = phase 5) presented in *H*.

Each base pair change in the nucleosome phasing changes the trajectory of the chromatin fiber by approximately 36° (1/10 of a full turn). To extend the model to include two adjacent TRs, we joined pairs of these single TR models at each of the 9 nucleosome phases to produce the complete set of 81 2TR models (9 phases for each tetrasome) (Fig. 8B and F), keeping the repeat length at 801 bp between LBS1 sites. For models with occupancy of only LBS1 and LBS2, we eliminated 11 of the 81 combinations from further consideration based on steric interference, either between two tetrasomes or the trajectory of connecting DNA (shown in purple and blue, Fig. 8B). For the models with full occupancy of three LBS sites, 50 models were ruled out (shown in purple and blue, Fig. 8F). Examples of 2TR phase combinations with or without steric hindrance are depicted in Fig. 8. The impact of both LANA-dependent DNA bending and nucleosome translational positioning on the construction of these 2TR models, and the ability of these parameters to eliminate certain tetrasome configurations, demonstrates the importance of these two factors for tether architecture.

### A Simple Molecular Model Captures Major Properties of 2TR LANA Tethers

Remarkably, our 2TR models captured key features of the dSTORM images. In particular, the model with full LANA occupancy of all three LBSs on both TRs and the second tetrasome joined to the DNA at “phase 8” mimicked features of the image (Fig. 5A, and see also Fig. 8E). The addition of three LANA dimers to each individual TR of the 2TR model produced an elongated arrangement of epitope binding sites and predicted the localization of their associated dye emissions. Furthermore, the long axes joining the epitope binding sites on each TR give rise to approximately a 45° angle between them by the combination of tetranucleosome positioning and LBS DNA bending. The distance between the centers of the two sets of epitopes was approximately 151 nm. Both the angle and spacing of emissions in the model approximated those of the experimental dSTORM data (Fig. 4B).

**Fig. 5.**
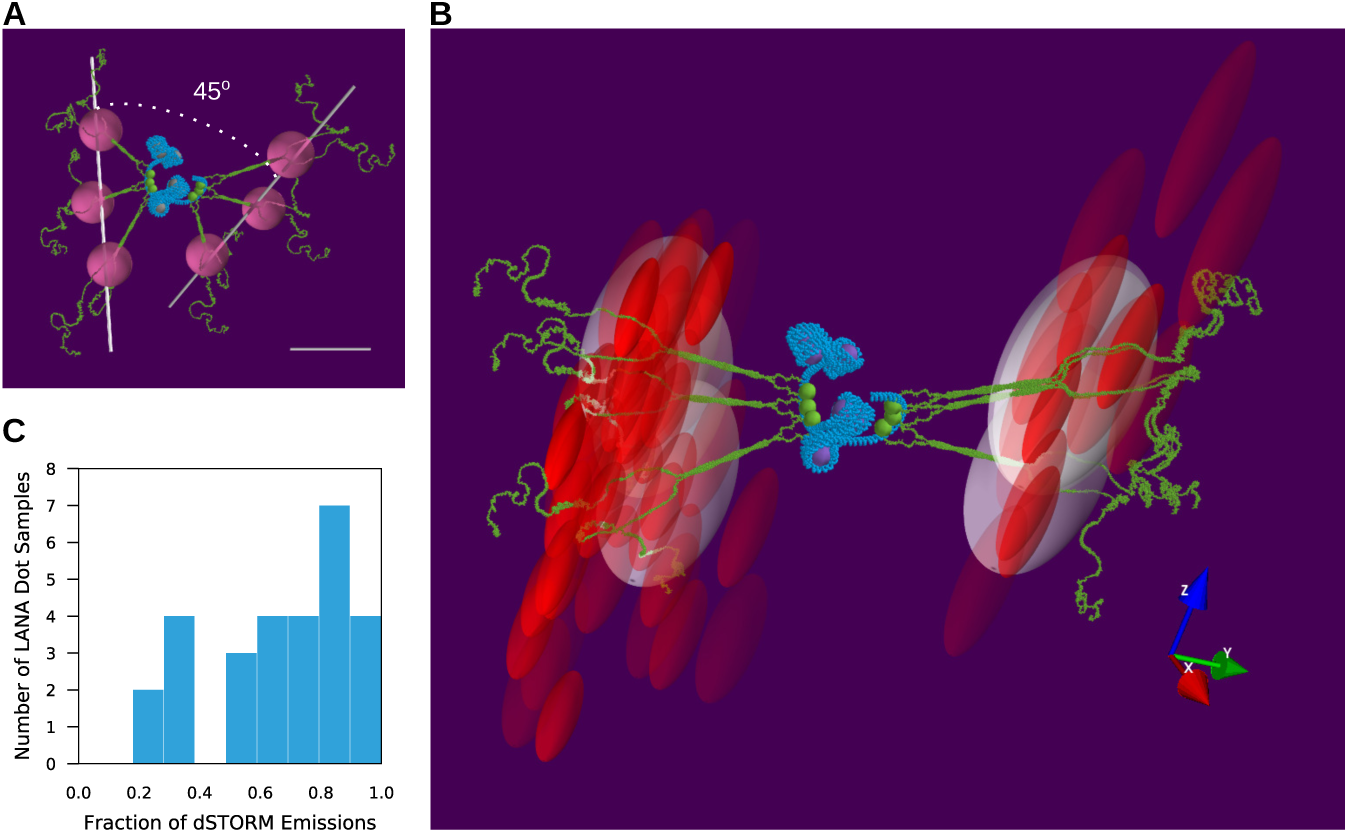
dSTORM analysis of p2TR tethers is consistent with a model featuring full LBS and nucleosome occupancy on two sequential TRs. (*A*) Model depicting two TRs (blue), each containing four nucleosomes (grey) and bound by three LANA dimers (green) occupying LBSs (scale bar = 50 nm). Predicted coiled-coil portions of LANA dimers extend outward at approximately right angles to the LBSs. Magenta spheres depict the position of a 13 nm-long anti-LANA mAb. Two white lines represent the best linear fit connecting three LANA dimers for each TR, forming a 45° angle. (*B*) Data from one p2TR example aligned with the model in (*A*). Red ellipsoids depict the probability volume of individual fluorophore emissions, scaled by their lateral and axial localization precision. White ellipsoids depict the model-predicted probability volumes scaled by data-derived localization precision. (*C*) Histogram displaying the distribution of the fraction of observed emissions whose localization precision volumes intersect with the model-predicted volume.

We tested the ability of our chromatin models to account for the dSTORM data of individual tethers by optimizing the fit of the modeled epitope sites (dye locations) to the experimental p2TR emissions data (Materials and Methods). We then determined the best fit models by scoring each by the fraction of dSTORM emissions that they captured. One example of such a fit is shown in Fig. 5B and Movie S1.

In a pair-wise comparison of models with LANA dimers at all three LBS sites versus models with LANA dimers only at LBS1 and LBS2, the full occupancy models were significantly better at accounting for the 28 p2TR datasets (*P* < 3.0 × 10^−12^, Paired T-test). Overall, the median fraction of dSTORM emissions accounted for by the optimal full occupancy models was 0.72, 95% CI [0.61, 0.83] (Fig. 5C) for the p2TR structures assembled in BJAB cells. We carried out the same analysis of p2TR LANA tethers expressed in COS-7 cells and found the median fraction of emissions covered to be 0.78, 95% CI [0.66, 0.94] (Fig. S3C and D). Interestingly, the optimal phasing between TRs was not the same for all 2TR tethers examined. Rather, this parameter varied from tether to tether, with phases 5-9 each contributing to the set of best fit models (Fig. 9). This range of phases suggests a basis for the shape variability seen among the individual LANA tethers comprised of larger numbers of TRs (Fig. 2).

**Fig. 9.**
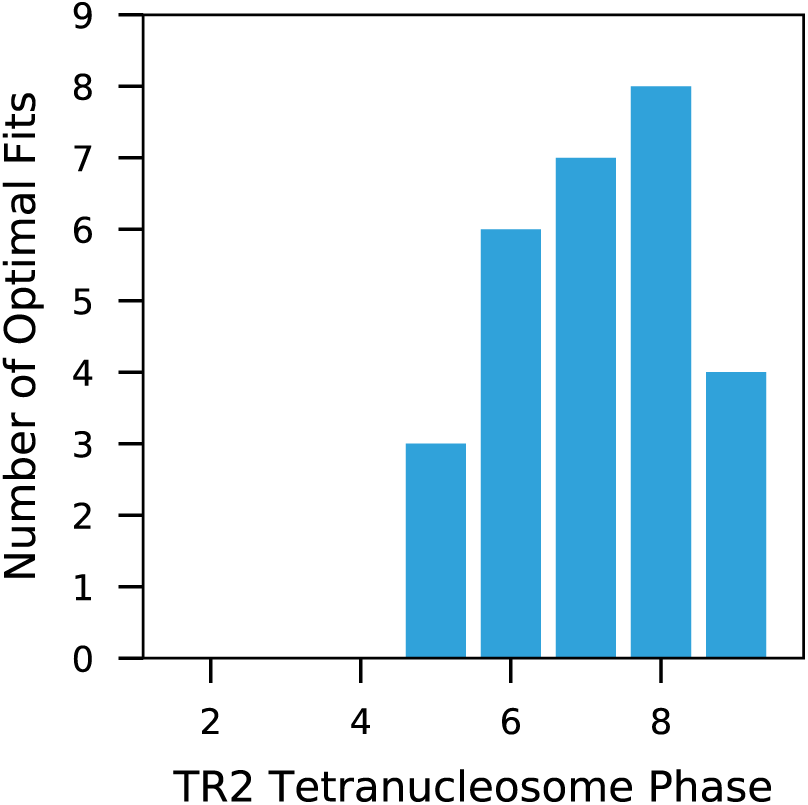
Multiple tetranucleosome translational positions are compatible with the p2TR dSTORM data in BJAB cells. The frequency of the occurrence of an optimal fit is plotted as a function of the tetranucleosome phasing of TR2 in models using 3 LANA dimers per TR.

## Discussion

Until now, LANA tethers have appeared as poorly defined nuclear punctae via epifluorescence microscopy. The localization data from this study offer new insights into the architecture of the tethers linking KSHV episomes to human chromatin. First, we found that the folding of tether chromatin was intrinsic to the viral episome and independent of the cellular environment. Consistent shapes and emission counts were measured for comparable TR episomes within the B cell-derived lines BCBL-1 and BJAB, the non-lymphocytic epithelial cell line SLKp, and even the monkey kidney fibroblast cell line COS-7. Second, regardless of the number, TRs were proportionally occupied by LANA dimers. This argues for a relatively uniform chromatin environment across the terminal repeats wherein LBS sites are equally accessible to LANA binding in all TRs. Third, we provided the first physical proof that TR chromatin is packaged with parameters that are characteristic of active chromatin. This may reflect the need for the TRs to provide alternate biological functions, such as the secure attachment of the episome to host nucleosomes versus the dynamic initiation of DNA replication. Fourth, the ability of dSTORM to resolve single TR units within 2TR clusters demonstrates that the episomal half of LANA dimers must be well ordered. If LANA dimers radiated randomly from the LBSs or were highly unstructured in their episomal half, we would be unable to separate discrete clusters in the analyses of plasmids containing 2 TRs. Finally, our data argue that all three LBS sites in individual TRs are likely to be fully occupied by LANA dimers. The discrete dimensions and positional asymmetry of the dye emission clusters in the 2TR data are not well accounted for if only two LBS sites are occupied, because the axis through the epitopes of each TR is shortened and there is less bending of the LBS DNA. Furthermore, models with three LANA dimers per TR consistently performed better than models with only two LANA dimers per TR in explaining dSTORM data for 26 independent 2TR datasets. The linear relationship between LANA dye emissions and TR number suggests that this is true for complete KSHV episomes as well.

We identified several key features of tether structure by modeling the 2TR data. First, the modeling revealed that the internal repeats of LANA likely comprise a coiled-coil domain for the LANA dimer. This domain was first predicted by in silico analysis, and then supported by quantitative analysis of LN53 antibody binding, and masks all but the two to three N-terminal most mAb binding sites. Second, the modeling highlighted an impact of translational positioning of the nucleosomes for each TR. Moving each tetranucleosome along the nine possible phases of the DNA helix alters the trajectory of the output TR DNA. This, in turn, greatly impacts the relative positions of the LBSs on adjacent TRs and, hence, their associated anti-LANA fluorophore emissions. The architecture driven by nucleosome positioning and LBS DNA bending positions the LANA N-terminal histone binding domains in a way that facilitates exploration of the surrounding environment, promoting their primary function of tethering to host chromatin.

While the 2TR model describes most of the tethers well, a few datasets are exceptions mainly due to the fact that they have relatively larger *R_g_* values. Others have proposed large multi-LANA dimer complexes based on crystallography of the C-terminal binding domain (10, 15, 38), and it is possible these 2TR exceptions reflect such assemblies. Further, our current model has some limitations. For example, it is unlikely that the fixed packing of tetranucleosomes we adapted from the crystallographic structure reflects the full range of conformations adapted by TR nucleosomes in vivo. Unlike the KSHV tether histones, the histones present in the tetranucleosome crystal structure lacked any posttranslational modifications. The presence of such modifications might alter the tetrasome structure, potentially loosening or tightening the histone packing, and thereby impact our model and its fit to our data set.

The current working model that we have developed captures major features of the KSHV tether, detailing the likely stoichiometry and relative positions of its major molecular components emanating from the TR region of the viral genome within the cellular milieu. The model also provides a platform for future experimentation that could include determining the nanoarchitecture of tethers with greater numbers of TRs. This approach has broad applicability to a wide variety of persistent viral pathogens, thus, potentially contributing to our ability to target them therapeutically.

## Materials and Methods

### Cell Lines

BCBL-1 cells have been in the laboratory of D. H.K. who was a co-author on the study first describing the line (45). BJAB cells were a gift from Don Ganem (Novartis Institutes for Biomedical Research) and are described by Menezes et al. (46). SLKp/Caki-1p (36) cells were a gift from Adam Grundhoff (Heinrich Pette Institute, Leibniz Institute for Experimental Virology, Hamburg, Germany) and Don Ganem and are described by Grundhoff and Ganem (35). SLKp cells were selected despite their inclusion in the ICLAC database of commonly misidentified cell lines because of their utility as an epithelial, non-lymphocytic line and their stable infection with BCBL-1-derived virus. iSLK-BAC16 cells were a gift from Rolf Renne (University of Florida, Department of Molecular Genetics and Microbiology) and are described by Brulois et al. (47). COS-7 cells were a gift from Jim Casanova (University of Virginia). BJAB and BCBL-1 cells had typical lymphocyte nonadherent morphology and were CD45 positive by immunofluorescence staining. BCBL-1 cells were syndecan-1 (CD138) positive by immunostaining and flow cytometry (26). BCBL-1, SLKp, and iSLK-BAC16 cells were LANA positive by immunofluorescence staining; BJAB and COS-7 cells were LANA negative.

Adherent cell lines were maintained in Dulbecco’s modified Eagle’s medium (Gibco) supplemented with 10% fetal bovine serum (Gibco). They were harvested from culture dishes with 0.05% trypsin after a wash with 1X phosphate buffered saline. Non-adherent cell lines were maintained in RPMI 1640 (Gibco) supplemented with 10% fetal bovine serum, 20 mM HEPES, 2 mM L-glutamine, 0.05 mM beta-mercaptoethanol, and 200 *μ*g/mL sodium bicarbonate. All experimental conditions for imaging took place in identical media but lacking phenol red. All cell lines were maintained at 37°C in 5% CO2 and tested negative for mycoplasma.

### Plasmids

The pcDNA3-LANA plasmid was a gift from Rolf Renne (University of Florida, Department of Molecular Genetics and Microbiology) and is described by Renne et al. (48). BAC16 was a gift from Jae Jung (University of Southern California) and is described by Brulois et al. (47). The p8TR plasmid is the 8TR-containing Z6 cosmid subcloned into modified pREP9, as described by Ballestas et al. (12). The plasmid, produced by the lab of Kenneth Kaye (Harvard Medical School), was a gift from Paul Lieberman (The Wistar Institute). Plasmids containing 0, 2, 5, and 7 terminal repeats were created by transforming DH5α E. coli (Thermo Fisher Scientific) with the p8TR plasmid and screening by *Pst*I restriction endonuclease digestion for colonies that showed recombination within the TR region and gave rise to a lower number of TRs. Insert size was confirmed by gel electrophoresis, as shown in Fig. S1A and B. The plasmids isolated from these E. coli were subsequently grown up in SURE E. coli (Agilent Technologies) to preserve fidelity of the new TR number.

### Antibodies

The anti-LANA antibody LN53 (rat, monoclonal) (49) was conjugated using an Alexa Fluor^®^ 647 succinimidyl ester kit (Thermo Fisher Scientific).

### Determining TR Number in BAC16 and BCBL-1 Virus

Linear viral DNA was obtained from BCBL-1 cells using the method described by Lagunoff et al. (16). BCBL-1-derived KSHV DNA and BAC16 DNA were digested with NotI + SpeI. The resulting digests were run on an agarose gel, which was then stained with ethidium bromide. We then plotted band intensity as a function of DNA length for each band present in a single copy per genome. We used this line to calculate the amount of DNA represented by the 801-bp band. For BAC16, this band gave a result of 16.2 kbp, which accounts for 20 TRs. Because the BAC16 is circular, one TR is not accounted for by this method, yielding a final calculation of 21 TRs. For the BCBL-1-derived viral DNA shown in Fig. S1E, the 801-bp band was representative of 36.0 kbp, which accounts for 45 TRs. Because this DNA is linear, two TRs are not accounted for by this method, yielding a final result of 47 TRs. For BCBL-1-derived KSHV, this experiment was performed in triplicate, and the resulting number of TRs was 43 +/−6.

### Transfections

1×10^7^ BJAB cells were transfected with 1*μ*g pNTR plasmid + 1*μ*g pcDNA3-LANA plasmid using Amaxa Nucleofector (Lonza) with Solution V on program T-020. Cells were harvested for analysis at 48 hours posttransfection.

### Infection of BJABs with BAC16 Virus

Infection of BJABs was performed as described by Plaisance-Bonstaff (48). In brief, iSLK-BAC16 cells were seeded at 3×10^5^/mL in a 6-well plate. After 24 hours, they were induced with doxycycline (1 *μ*g/mL) and sodium butyrate (1 mM). 48 hours later, the media was replaced with 3 mLs RPMI containing BJAB cells at a density of 3×10^5^/mL. Four days later, BJABs were removed from the iSLK-BAC16 cells and cultured for 14 days with hygromycin selection (100 *μ*g/mL) before staining. A separate, identically treated well of iSLK-BAC16 cells without BJABs was cultured for 7 days, at which point all induced iSLK-BAC16s were found to be dead by Trypan blue exclusion.

### Antibodies

The anti-LANA antibody LN53 (rat, monoclonal) (49) was conjugated using an Alexa Fluor^®^ 647 succinimidyl ester kit (Thermo Fisher Scientific). The approximate number of dye molecules per antibody (three) was calculated according to the manufacturer’s instructions.

### dSTORM Sample Preparation

Cells were fixed in 4% paraformaldehyde for 10 minutes. After a 20-minute rat IgG block at 4 *μ*g/mL, Alexa Fluor^®^-conjugated LN53 antibody was applied at a concentration of 1:400 in BD Perm/Wash (BD Biosciences) and incubated for 40 minutes. Cells were washed in Perm/Wash and DAPI was then applied at 0.5*μ*g/mL for 15 minutes. Coverslips (Cat. #72204-01, Electron Microscopy Sciences) were cleaned with alkaline peroxide. Their centers were masked off and TetraSpeck 0.1*μ*m beads (Thermo Fisher Scientific) were applied to the rest of the surface; these were attached to the slide at 100°C for 30 min. The center masks were removed and the coverslips were coated in poly-L-lysine for 1 hour. The poly-L-lysine solution was aspirated and the coverslips were air-dried overnight. Cells were deposited (Cytospin 2, Shandon) onto the centers of these prepared coverslips for 4 minutes at 500 rpm. Samples for dSTORM analysis were mounted in a solution of 0.22 mg/mL glucose oxidase, 40 *μ*g/mL catalase, 0.14M beta-mercaptoethanol, and 0.55M glucose.

### Instrumentation

dSTORM data were collected using custom instrumentation based on an Olympus IX81-ZDC2 inverted microscope (Olympus America Inc., Melville, NY, USA) configured in a standard geometry for localization microscopy (50, 51). The illumination beam from a 643 nm diode laser (DL-640-100, CrystaLaser, Reno, NV, USA), coupled to a laser clean-up filter (ZET640/20X, Chroma Technology Corp., Rockingham, VT, USA) and a 5X Galilean beam expander (BEO5M, Thorlabs, Inc., Newton, NJ, USA), was focused on the backplane of a 60X, 1.2NA, water-immersion objective (UPSLAPO60XW, Olympus) by means of a f+350 mm lens (Thorlabs) to yield an illumination profile with a 1/e^2^ radius of 34 *μ*m. Ground state depletion and dSTORM readout of Alexa Fluor^®^ 647 dye molecules was achieved with a laser intensity of 1.9 kW/cm^2^ as measured at the sample. Single molecule emissions were collected through the same objective using a filter set comprising a dichroic mirror (Di01-R635, Semrock, Rochester, NY, USA) and emission filter (FF01-692/40, Semrock). The image was expanded using f+75 mm and f+150 mm lenses (Thorlabs) arranged as a ~2X telescope and acquired with an EMCCD camera (iXonEM DU-897D-CS0-BV, Andor Technology PLC, South Windsor, CT, USA) to yield an image pixel size of 0.134 *μ*m (52). Astigmatism for axial localization was introduced using a f+1000 mm cylindrical lens (LJ1516RM-A, Thorlabs) placed in the acquisition path (53).

### dSTORM Data Acquisition

dSTORM images were acquired using a water-immersion objective to minimize axial compression of the data for LANA tethers located far from the coverslip (54). Because of the sensitivity of water-immersion lenses to the tilt of the coverslip (55), axial localization was calibrated for each sample by imaging pointsource fluorescent beads attached to the same coverslip as the samples being studied. Images of the astigmatic point spread function were acquired every 50 nm in the axial dimension, spanning 2 *μ*m across the focal plane, controlled by a 3-axis nanopositioning stage (Nano-LPS100, Mad City Labs Inc., Madison, WI, USA). dSTORM images were collected at a frame rate of 32.4 Hz, for a total of 15,000 to 25,000 frames, using an EM gain of 100 and pre-amplifier gain of 5X on the iXon camera. Transmitted light images were collected every 200 frames to provide subsequent correction for lateral drift during analysis, while axial positioning was stabilized by the ZDC2 instrumentation of the microscope. An epifluorescence image of DAPI staining was saved after the dSTORM data acquisition and the few nuclei exhibiting chromosome condensation were excluded from further analysis.

### dSTORM Data Processing

dSTORM images were processed using a pipeline of custom python modules, implementing standard single molecule localization algorithms (51). In-house software tools utilized `numpy` (version 1.11.2) and `scipy` (version 0.18.1) scientific computing libraries (56, 57). Candidate dye emissions were identified by size and photon threshold criteria and localized within an 11 x 11 pixel region of interest. Each point spread function (PSF) was evaluated by fitting with a two-dimensional Gaussian ellipse by least squares to estimate parameters for the x and y center coordinates, amplitude, major and minor axes radii, and an offset. The lateral localization precision was calculated for each emission as described by Thompson et al. (52). For each emission, photon yield was calculated as the product of the Gaussian amplitude and area, while the background pixel noise was estimated as the standard deviation of intensities from the perimeter pixels of the region of interest. The axial location of a molecule was determined using its major and minor axis radii from the 2D Gaussian fit to interrogate the standard curve established from the point source fluorescent beads (53, 58) and its axial localization precision was calculated as described by DeSantis et al. (58). The transmitted light images obtained through dSTORM acquisition were used to determine lateral image drift using a sub-pixel crosscorrelation algorithm (59) and dSTORM molecule coordinates were corrected by interpolation.

### Determination of *R_g_* and Statistical Treatments

LANA tethers were characterized by their radius of gyration, calculated as the root mean squared vectors of the emissions using the standard representation,

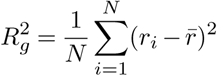

where *r_i_* is the location of individual emissions and *r̅* is the centroid of all the emissions in the tether. The coordinates of *r̅* were calculated as the average of the coordinates for all emissions weighted by the inverse of their respective lateral and axial localization precision.

Summary and comparative statistics were based on nonparametric approaches. The 95% confidence intervals for median estimates were calculated by bootstrap resampling. The similarity of the BCBL-1 and SLKp tethers were evaluated by Mood’s Median test (*α* = 0.05). Power curve analysis by bootstrap resampling of the BCBL-1 (N=65) and SLKp (N=34) populations revealed that, with a statistical power of 0.8, Mood’s Median was expected to reject the null hypothesis of equality if the SLKp tether median *R_g_* values were ≤ 68 nm (20%) smaller or ≥ 64 nm (19%) larger than BCBL-1.

### LANA Tether Analysis

The software tools used to analyze and visualize the LANA tethers were supported by the additional Python libraries `scikit-learn` (version 0.17.1) (60), `statsmodels` (version 0.6.1) (61), Geometric Tools (version wm5) (62), `matplotlib` (version 1.5.3) (63), and `mayavi.mlab` (version 4.5.1) (64). The dSTORM signals for individual LANA tethers were isolated from the data sets and further processed for analysis. The axial localization precision for each emission was re-evaluated by calculating the individual axial position 1000 times using values of the major and minor PSF widths randomly drawn from their respective error distributions. The larger of this Monte Carlo estimate and the original analytical calculation was saved as the working axial localization precision and emissions with values greater than ±60 nm were discarded.

To determine aggregate parameters individual LANA tethers were uniformly oriented by geometry-based principle component analysis and aligned at their centers of mass. The dimensions of the aggregate structures along the X, Y, and Z axes were estimated as the coordinates spanning quantile probabilities from 2.5% to 97.5% of the distribution along each axis. Solid 3D representations of tether emissions are depicted by spheres with a radius equal to their median axial localization precision. Similarly, tether images are rendered as Gaussian spots scaled to a width that is twice the median axial localization precision.

Individual 2TR LANA tethers were analyzed further by dividing their emissions into two groups using k-means clustering. The midpoint between the centers of the two clusters was translated to the zero origin and the complete tether was then oriented to place the two cluster centers along the x-axis on either side of the origin. The tether was then rotated about the x-axis to achieve the greatest variance in the y-dimension. Vectors fit through the long dimensions of the two clusters defined a “V” geometry, and each tether was then reflected horizontally or vertically or both such that the wider separation was oriented towards the positive y-axis and greatest tilt from vertical was oriented towards the positive x-axis. The complete set of 2TR LANA tethers were then superimposed at their origins to evaluate the parameters of the aggregate structure.

### Modeling 2TR LANA tethers

The software written to model 2TR LANA tethers utilized additional python modules from the programming libraries `Biopython` (version 1.68) (65, 66) and `PeptideBuilder` (version 1.0.1) (67), and the application `PyMOL` (version 1.7.6) (68).

The X-ray crystal structure of a dimer of LANA C-terminal DNA binding domains in complex with LBS1 DNA suggests that the upstream LANA peptides may enter on either side of the episomal DNA (Fig. 6B) (15) (PDB 4UZB). Therefore, in modeling LANA dimers, we placed the LN53 antibody epitopes on the N-terminal side of a coiled-coil domain which is then connected by a domain of unknown structure to the dimer of C-terminal DNA binding domains. This model is similar to a more general outline proposed previously (15). We modeled domains of unknown structure based on psi and phi angles from their predicted secondary structure. These were done solely to provide plausible constraints on the placement of the EQEQ epitopes relative to the LBS, and with the exception of the coiled-coil model (69) and C-terminal DNA binding domains with known crystal structures, the models are not expected to match actual LANA protein folding.

**Fig. 6.**
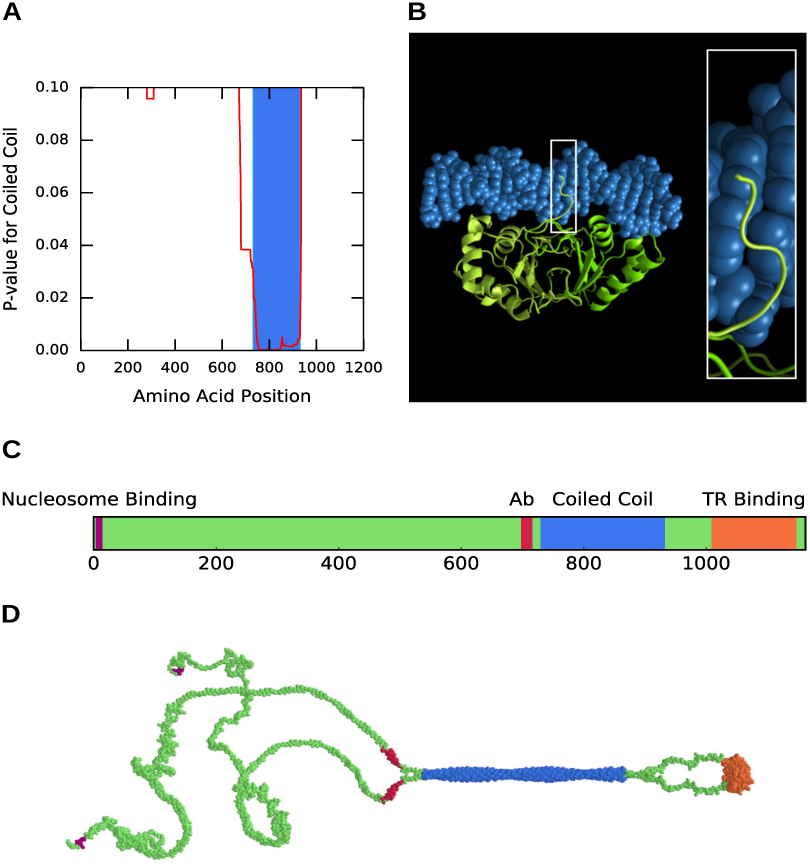
Modeling LANA dimers. (*A*) `PairCoil2` (40) predicts a coiled-coil domain in LANA. The position of the domain is shown as a plot of probability versus amino acid number (LANA sequence GenBank: U75698.1). (*B*) The structure of a LANA dimer bound to LBS1, adapted from (15). Shown is the wrapping of LANA around the DNA to the side opposite LBS binding (inset). (*C*) Diagram of the various domains of LANA, including the nucleosome binding domain (purple), LN53 Ab binding sites outside the coiled coil region (red), coiled-coil region (blue), and TR binding domain (orange). LN53 Ab sites C-terminal to the red region were discounted from the TR modeling due to their being buried within the coiled-coil. (*D*) Structural rendering of LANA, showing the features and color scheme as in *C*.

Complete sets of 2TR models were constructed in which the TR1 and TR2 tetrasomes were arranged over 9 translational positions each (Fig. S6). The critical parameter for fitting the models to the dSTORM data is the phase of the tetranucleosome in TR2 because it is located between the two sets of LBS DNA sequences. While the phase of the tetranucleosome in TR1 is presumably important for the connection to flanking chromatin, its phase does not affect the structural relationship between the LANA dimers in the two TRs. Therefore, for the 2-dimer models, we fit each of the 28 p2TR datasets with the 9 chromatin models that combined TR1 phase 3 with TR2 phases 1 through 9 (Fig. S6B). For the 3-dimer models, we fit the same 28 p2TR datasets with the 6 chromatin models that combined TR1 phase 6 with TR2 phases 4 through 9 (Fig. S6F).

2TR models were fit to the dSTORM data using a limited memory approximation of the Broyden-Fletcher-Goldfarb-Shanno algorithm for bound constrained minimization (70) as implemented in `SciPy`. Each of the two clusters of dSTORM emissions for individual tethers was further divided into either two or three clusters by k-means to yield either four or six centroids per tether, for the 2-dimer and three-dimer models respectively. To begin optimization, the center of mass of the epitope binding sites in a model was aligned with the global center of mass of the cluster centroids in the dSTORM data. The initial rotational orientation of the model was randomized. Eighteen parameters were then varied to minimize the Euclidean distance between the model epitope sites and their respective dSTORM cluster centers. For fitting, the model was allowed three dimensions of rotation and three dimensions of translation. The four or six epitope sites in the models were treated as linear vectors of a fixed 51 nm in length originating at the LBS sites and they were allowed to rotate freely but constrained to a maximum 45° tilt, or less, from perpendicular to the DNA. The best fit for each model was then evaluated for its ability to account for the dSTORM emissions. The Gaussian probability volumes of the dSTORM emissions were scaled by 2.2 times their lateral and axial localization precision. Similarly, the model epitope site probability volumes were scaled by 2.2 times the median axial and lateral localizations of their associated dSTORM emissions. The fit was then scored for the fraction of the dSTORM ellipsoids intersecting with the epitope ellipsoids of the model.

## ACKNOWLEDGEMENTS

We thank Barbara Knowles, Samuel Hess, Travis Gould, and Mudalige Gunewardene for an introduction to super-resolution microscopy and advice on instrumentation, and Stefan Bekiranov for discussions. This work was supported in part by the University of Virginia Cancer Center P30CA044579 (D.H.K., M.M.S.). D.H.K. acknowledges support of the National Institute of Dental and Craniofacial Research R01DE022291. M.M.S. acknowledges support of the National Institute of General Medical Sciences RC1GM091175 and R01GM116994.

---

**Video S1. Animation of the three-dimensional modeling of LANA binding to TR DNA and the resulting Ab-bound fluorophore emissions.** Each of the many fluorophore emissions captured by dSTORM from a single 2TR example is presented as a red ellipsoid encompassing that emission’s median three-dimensional localization precision. These data are superimposed on the model of 2TRs presented in Figure 5A and B. Emerging magenta spheres represent the 13 nm radius encompassing the potential localization of the bound anti-LANA mAb, as in Figure 5B. The video zooms in to more clearly show the architecture surrounding the TR DNA, LANA C-terminal LBS binding, and associated tetrasomes. Finally, white ellipsoids appear, denoting model-predicted fluorophore emissions at one standard deviation of the localization precision.

**Fig. S1.**
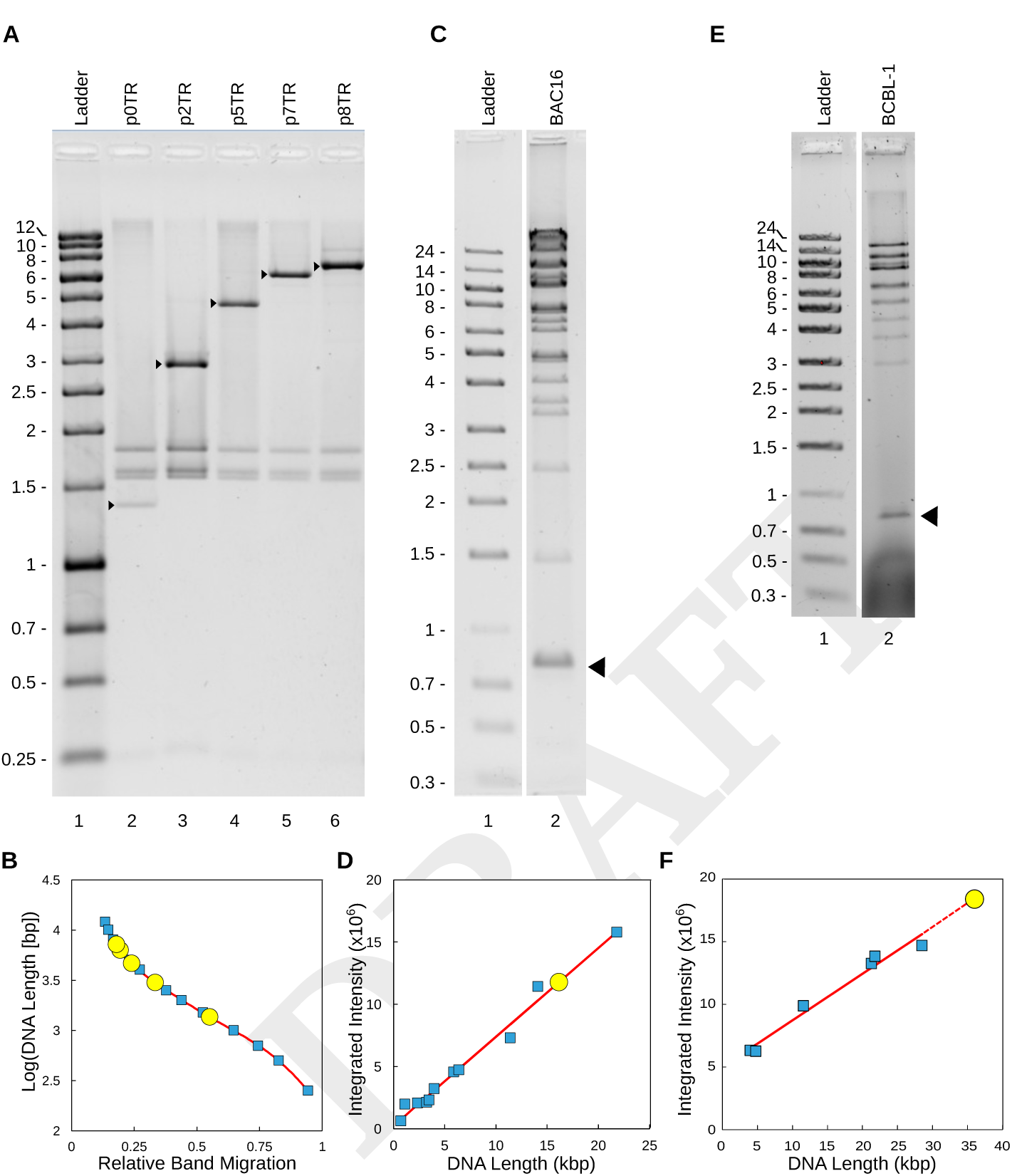
Number of TRs in each pNTR plasmid and BAC16. (*A*) Plasmids with variable (*N*) numbers of TRs (pNTR) were digested with PstI, which cuts on either side of the TR-containing insert. Length of the released fragment (arrowheads) allowed determination of TR number. Lane 1, ladder (kbp), lane 2, plasmid lacking TRs, lanes 3-6, plasmids with 2, 5, 7, and 8 TRs, respectively. (*B*) Relative mobilities of DNA standards (blue squares) were used to construct a best-fit curve for the log of DNA length as a function of migration (red line). Lengths of PstI fragments from plasmids with variable numbers of TRs (yellow circles) were determined from the standard curve based on their migration. (*C*) Lane 1, ladder (kbp). Lane 2, Notl + Spel digest of BAC16 isolating the 801-bp TR fragments (arrowhead). (*D*) The number of TRs within BAC16 was determined from the intensity of ethidium bromide staining. Fragments present once per BAC (blue squares) were used to determine a best-fit line (red) for band intensity as a function of DNA length (*R*^2^ = 0.98). Intensity of the TR-containing fragment (yellow circle) was then mapped to this standard curve, yielding a calculation of 21 TRs per BAC16 genome. (*E*) Lane 1, ladder (kbp). Lane 2, Notl + Spel digest of BCBL-1-derived KSHV DNA, isolating the 801-bp TR fragments (arrowhead). (*F*) The number of TRs in BCBL-1-derived virus was determined as in D and was calculated to be 45 (*R*^2^ = 0.97) for the experiment shown. This experiment was performed in triplicate, yielding a final result of 43 +/− 6 TRs per BCBL-1 episome.

**Fig. S2.**
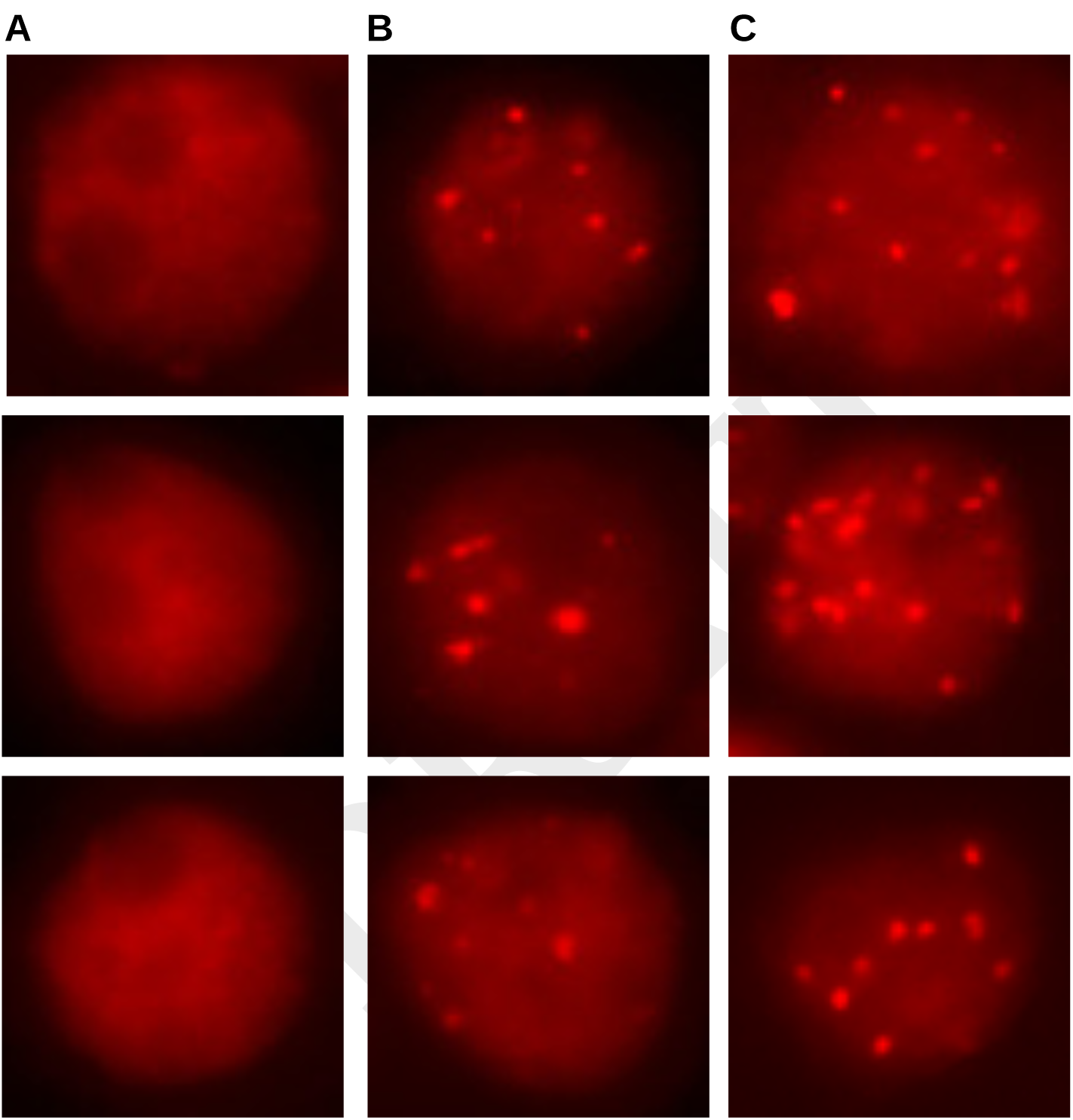
Punctate LANA tethers are visible only in the presence of TRs. (*A-B*) Epifluorescence images show three representative BJAB cells transfected with LANA-expressing plasmid plus either (*A*) p0TR plasmid, resulting in diffuse, non-punctate LANA staining, or (*B*) p8TR plasmid, resulting in punctate LANA staining. (*C*) BJAB cells co-cultured with induced iSLK-BAC16 cells also resulted in punctate LANA staining.

**Fig. S3.**
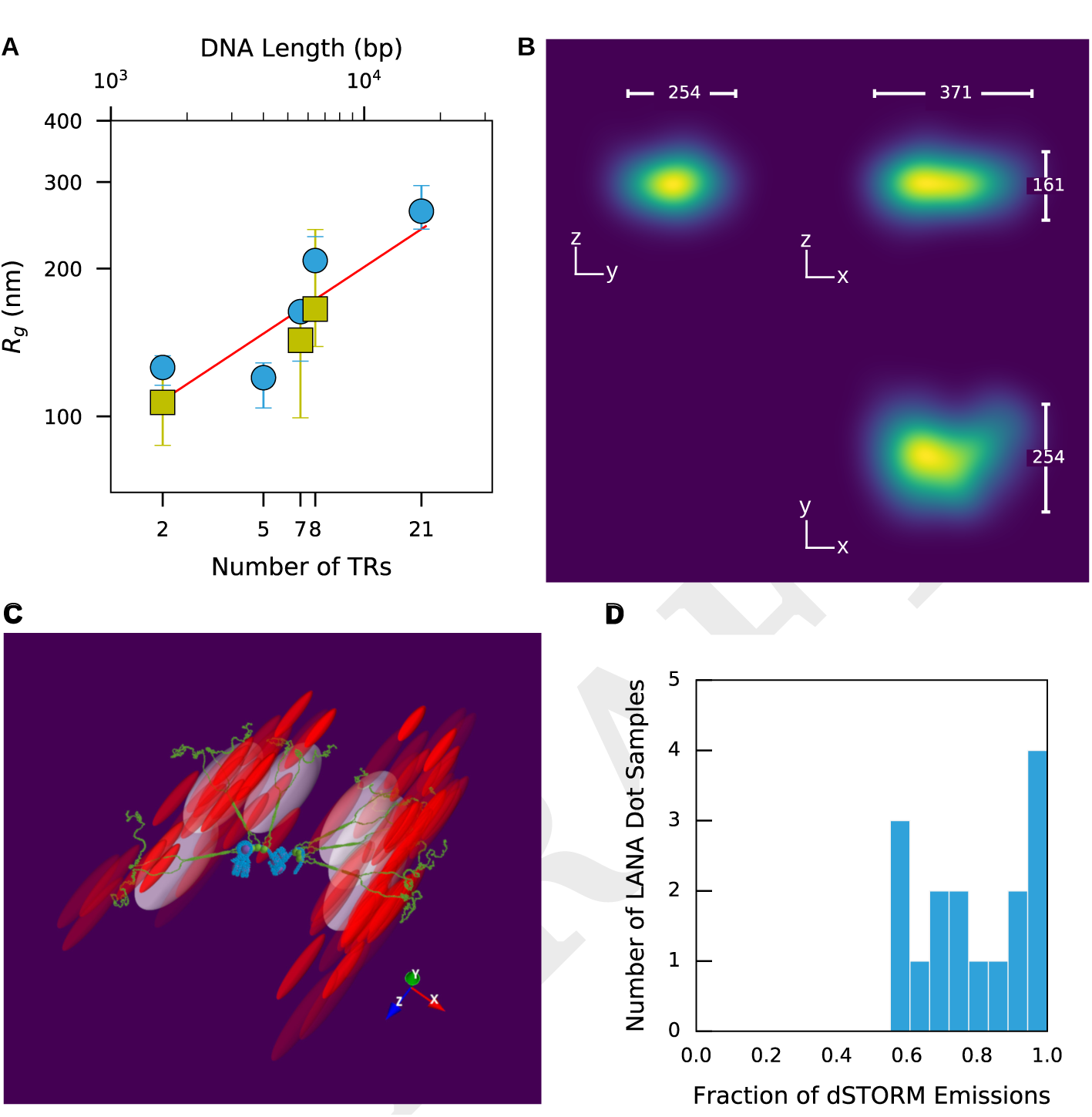
Analysis of LANA tethers in COS-7 cells. (*A*) Scaling of *R_g_* with increasing number of TRs in COS-7 cells supports the results seen in BJAB cells. A plot of *R_g_* vs TR number for p2TR, p7TR, and p8TR in COS-7 (yellow squares), shown in comparison with the BJAB data (blue circles). The combined data produce a scaling exponent of c = 0.35 ± 0.07, *R*^2^ = 0.89. (*B*) COS-7 p2TR data shows two clusters of emissions for each tether, similar to that seen in BJAB cells. Compilation of emission data from 16 p2TR tethers in COS-7 cells; emissions are rendered as in Fig. 2. Median dimensions (nm) of the combined data are shown. (*C*) Data from one p2TR example aligned with a model similar to that shown in Fig. 5A. Red ellipsoids depict the probability volume of individual fluorophore emissions, scaled by their lateral and axial localization precision. White ellipsoids depict the model-predicted probability volumes scaled by data-derived localization precision. (*D*) Histogram displaying the distribution of the fraction of observed emissions whose localization precision volumes intersect with the model-predicted volume. Median overlap fraction = 0.78, 95% CI [0.66, 0.94].

## Bibliography

1. Y Chang, E Cesarman, M. Pessin, F Lee, J Culpepper, D. Knowles, and P. Moore. Identification of herpesvirus-like DNA sequences in AIDS-associated Kaposi’s sarcoma. Science, 266(5192):1865–1869, December 1994. ISSN 0036-8075, 1095-9203. doi: 10.1126/science.7997879.

2. Ethel Cesarman, Yuan Chang, Patrick S. Moore, Jonathan W. Said, and Daniel M. Knowles. Kaposi’s Sarcoma-Associated Herpesvirus-Like DNA Sequences in AIDS-Related Body-Cavity-Based Lymphomas. New England Journal of Medicine, 332(18):1186–1191, May 1995. ISSN 0028-4793, 1533-4406. doi: 10.1056/NEJM199505043321802.

3. Jean Soulier, Laurence Grollet, Eric Oksenhendler, Patrice Cacoub, Dominique Cazals-Hatem, Paul Babinet, M. F. d’Agay, Jean-Pierre Clauvel, Martine Raphael, Laurent Degos, and others. Kaposi’s sarcoma-associated herpesvirus-like DNA sequences in multicentric Castleman’s disease [see comments]. Blood, 86(4):1276–1280, 1995.

4. Dirk Dittmer, Michael Lagunoff, Rolf Renne, Katherine Staskus, Ashley Haase, and Don Ganem. A Cluster of Latently Expressed Genes in Kaposi’s Sarcoma-Associated Herpesvirus. J Virol, 72(10):8309–8315, October 1998. ISSN 0022-538X.

5. Dean H. Kedes, Michael Lagunoff, Rolf Renne, and Don Ganem. Identification of the gene encoding the major latency-associated nuclear antigen of the Kaposi’s sarcoma-associated herpesvirus. Journal of Clinical lnvestlgation, 100(10):2606,1997.

6. L. Rainbow, G. M. Platt, G. R. Simpson, R. Sarid, S. J. Gao, H. Stoiber, C. S. Herrington, P. S. Moore, and T. F. Schulz. The 222- to 234-kilodalton latent nuclear protein (LNA) of Kaposi’s sarcoma-associated herpesvirus (human herpesvirus 8) is encoded by orf73 and is a component of the latency-associated nuclear antigen. J. Virol., 71(8):5915–5921, August 1997. ISSN 0022-538X, 1098-5514.

7. A. J. Barbera. The Nucleosomal Surface as a Docking Station for Kaposi’s Sarcoma Herpesvirus LANA. Science, 311(5762):856–861, February 2006. ISSN 0036-8075, 1095-9203. doi: 10.1126/science.1120541.

8. Andrew J. Barbera, Jayanth V. Chodaparambil, Brenna Kelley-Clarke, Karolin Luger, and Kenneth M. Kaye. Kaposi’s sarcoma-associated herpesvirus LANA hitches a ride on the chromosome. Cell Cycle, 5(10):1048–1052, 2006.

9. J. You, V. Srinivasan, G. V. Denis, W. J. Harrington, M. E. Ballestas, K. M. Kaye, and P. M. Howley Kaposi’s Sarcoma-Associated Herpesvirus Latency-Associated Nuclear Antigen Interacts with Bromodomain Protein Brd4 on Host Mitotic Chromosomes. Journal of Virology, 80(18):8909–8919, September 2006. ISSN 0022-538X. doi: 10.1128/JVI.00502-06.

10. Jan Hellert, Magdalena Weidner-Glunde, Joern Krausze, Ulrike Richter, Heiko Adler, Roman Fedorov, Marcel Pietrek, Jessica Rückert, Christiane Ritter, Thomas F. Schulz, and others. A structural basis for BRD2/4-mediated host chromatin interaction and oligomer assembly of Kaposi sarcoma-associated herpesvirus and murine gammaherpesvirus LANA proteins. PLoS Pathog, 9(10):e1003640, 2013.

11. James J. Russo, Roy A. Bohenzky, Ming-Cheng Chien, Jing Chen, Ming Yan, Dawn Maddalena, J. Preston Parry, Daniela Peruzzi, Isidore S. Edelman, Yuan Chang, and Patrick S. Moore. Nucleotide sequence of the Kaposi sarcoma-associated herpesvirus (HHV8). Proc NatlAcadSciUSA, 93(25):14862–14867, December 1996.

12. M. E. Ballestas. Efficient Persistence of Extrachromosomal KSHV DNA Mediated by Latency-Associated Nuclear Antigen. Science, 284(5414):641–644, April 1999. ISSN 00368075,10959203. doi: 10.1126/science.284.5414.641.

13. A. C. Garber, M. A. Shu, J. Hu, and R. Renne. DNA Binding and Modulation of Gene Expression by the Latency-Associated Nuclear Antigen of Kaposi’s Sarcoma-Associated Herpesvirus. Journal of Virology, 75(17):7882–7892, September 2001. ISSN 0022-538X. doi: 10.1128/JVI.75.17.7882-7892.2001.

14. A. C. Garber, J. Hu, and R. Renne. Latency-associated Nuclear Antigen (LANA) Cooperatively Binds to Two Sites within the Terminal Repeat, and Both Sites Contribute to the Ability of LANA to Suppress Transcription and to Facilitate DNA Replication. Journal of BiologicalChemistry, 277(30):27401–27411, July 2002. ISSN 0021-9258, 1083-351X. doi: 10.1074/jbc.M203489200.

15. Jan Hellert, Magdalena Weidner-Glunde, Joern Krausze, Heinrich Lünsdorf, Christiane Ritter, Thomas F. Schulz, and Thorsten Lührs. The 3d structure of Kaposi sarcoma herpesvirus LANA C-terminal domain bound to DNA. Proceedings of the National Academy of Sciences, 112(21):6694–6699, May 2015. ISSN 0027-8424, 1091-6490. doi: 10.1073/pnas.1421804112.

16. Michael Lagunoff and Don Ganem. The Structure and Coding Organization of the Genomic Termini of Kaposi’s Sarcoma-Associated Herpesvirus (Human Herpesvirus 8). Virology, 236(1):147–154, September 1997. ISSN 00426822. doi: 10.1006/viro.1997.8713.

17. Jean-Gabriel Judde, Vincent Lacoste, Josette Brière, Eric Kassa-Kelembho, Emmanuel Clyti, Pierre Couppié, Carmen Buchrieser, Micheline Tulliez, Jacques Morvan, and Antoine Gessain. Monoclonality or Oligoclonality of Human Herpesvirus 8 Terminal Repeat Sequences in Kaposi’s Sarcoma and Other Diseases. J Natl Cancer Inst, 92(9):729–736, May 2000. ISSN 0027-8874. doi: 10.1093/jnci/92.9.729.

18. Mary E. Ballestas and Kenneth M. Kaye. Kaposi’s Sarcoma-Associated Herpesvirus Latency-Associated Nuclear Antigen 1 Mediates Episome Persistence through cis-Acting Terminal Repeat (TR) Sequence and Specifically Binds TR DNA. J. Virol., 75(7):3250–3258, April 2001. ISSN 0022-538X, 1098-5514. doi: 10.1128/JVI.75.7.3250-3258.2001.

19. Takashi Komatsu, Mary E. Ballestas, Andrew J. Barbera, Brenna Kelley-Clarke, and Kenneth M. Kaye. KSHV LANA1 binds DNA as an oligomer and residues N-terminal to the oligomerization domain are essential for DNA binding, replication, and episome persistence. Virology, 319(2):225–236, February 2004. ISSN 0042-6822. doi: 10.1016/j.virol.2003.11. 002.

20. Feng-Chun Ye, Fu-Chun Zhou, Seung Min Yoo, Jian-Ping Xie, Philip J. Browning, and Shou-Jiang Gao. Disruption of Kaposi’s Sarcoma-Associated Herpesvirus Latent Nuclear Antigen Leads to Abortive Episome Persistence. J. Virol., 78(20):11121–11129, October 2004. ISSN 0022-538X, 1098-5514. doi: 10.1128/JVI.78.20.11121-11129.2004.

21. Franceline Juillard, Min Tan, Shijun Li, and Kenneth M. Kaye. Kaposi’s Sarcoma Herpesvirus Genome Persistence. Frontiers in Microbiology, 7, August 2016. ISSN 1664-302X. doi: 10.3389/fmicb.2016.01149.

22. S. J. Gao, L. Kingsley, M. Li, W. Zheng, C. Parravicini, J. Ziegler, R. Newton, C. R. Rinaldo, A. Saah, J. Phair, R. Detels, Y. Chang, and P. S. Moore. KSHV antibodies among Americans, Italians and Ugandans with and without Kaposi’s sarcoma. Nat. Med., 2(8):925–928, August 1996. ISSN 1078-8956.

23. D. H. Kedes, E. Operskalski, M. Busch, R. Kohn, J. Flood, and D. Ganem. The seroepidemiology of human herpesvirus 8 (Kaposi’s sarcoma-associated herpesvirus): distribution of infection in KS risk groups and evidence for sexual transmission. Nat. Med., 2(8):918–924, August 1996. ISSN 1078-8956.

24. P. S. Moore, S. J. Gao, G. Dominguez, E. Cesarman, O. Lungu, D. M. Knowles, R. Garber, P. E. Pellett, D. J. McGeoch, and Y. Chang. Primary characterization of a herpesvirus agent associated with Kaposi’s sarcomae. J. Virol., 70(1):549–558, January 1996. ISSN 0022-538X.

25. G. R. Simpson, T. F. Schulz, D. Whitby, P. M. Cook, C. Boshoff, L. Rainbow, M. R. Howard, S. J. Gao, R. A. Bohenzky, P. Simmonds, C. Lee, A. de Ruiter, A. Hatzakis, R. S. Tedder, I. V. Weller, R. A. Weiss, and P. S. Moore. Prevalence of Kaposi’s sarcoma associated herpesvirus infection measured by antibodies to recombinant capsid protein and latent immunofluorescence antigen. Lancet, 348(9035):1133–1138, October 1996. ISSN 0140-6736. doi: 10.1016/S0140-6736(96)07560-5.

26. L. A. Adang, C. H. Parsons, and D. H. Kedes. Asynchronous Progression through the Lytic Cascade and Variations in Intracellular Viral Loads Revealed by High-Throughput Single-Cell Analysis of Kaposi’s Sarcoma-Associated Herpesvirus Infection. Journal of Virology, 80(20):10073–10082, October 2006. ISSN 0022-538X. doi: 10.1128/JVI.01156-06.

27. Murray A. Cotter and Erle S. Robertson. The Latency-Associated Nuclear Antigen Tethers the Kaposi’s Sarcoma-Associated Herpesvirus Genome to Host Chromosomes in Body Cavity-Based Lymphoma Cells. Virology, 264(2):254–264, November 1999. ISSN 00426822. doi: 10.1006/viro.1999.9999.

28. Brenna Kelley-Clarke, Mary E. Ballestas, Takashi Komatsu, and Kenneth M. Kaye. Kaposi’s sarcoma herpesvirus C-terminal LANA concentrates at pericentromeric and peri-telomeric regions of a subset of mitotic chromosomes. Virology, 357(2):149–157, January 2007. ISSN 0042-6822. doi: 10.1016/j.virol.2006.07.052.

29. Thomas Günther and Adam Grundhoff. The Epigenetic Landscape of Latent Kaposi Sarcoma-Associated Herpesvirus Genomes. PLoS Pathogens, 6(6):e1000935, June 2010. ISSN 1553-7374. doi: 10.1371/journal.ppat.1000935.

30. Shuhei Sakakibara, Keiji Ueda, Ken Nishimura, Eunju Do, Eriko Ohsaki, Toshiomi Okuno, and Koichi Yamanishi. Accumulation of Heterochromatin Components on the Terminal Repeat Sequence of Kaposi’s Sarcoma-Associated Herpesvirus Mediated by the Latency-Associated Nuclear Antigen. J. Virol., 78(14):7299–7310, July 2004. ISSN 0022-538X, 1098-5514. doi: 10.1128/JVI.78.14.7299-7310.2004.

31. W. Stedman, Z. Deng, F. Lu, and P. M. Lieberman. ORC, MCM, and Histone Hyperacetylation at the Kaposi’s Sarcoma-Associated Herpesvirus Latent Replication Origin. Journal of Virology, 78(22):12566–12575, November 2004. ISSN 0022-538X. doi: 10.1128/JVI.78.22.12566-12575.2004.

32. Rui Sun, Xiaohua Tan, Xing Wang, Xiaodong Wang, Lei Yang, Erle S. Robertson, and Ke Lan. Epigenetic Landscape of Kaposi’s Sarcoma-Associated Herpesvirus Genome in Classic Kaposi’s Sarcoma Tissues. PLOS Pathogens, 13(1):e1006167, January 2017. ISSN 1553-7374. doi: 10.1371/journal.ppat.1006167.

33. Magdalena Weidner-Glunde, Giuseppe Mariggiò, and Thomas F. Schulz. Kaposi’s Sarcoma Herpesvirus Latency-Associated Nuclear Antigen (LANA): Replicating and Shielding Viral DNA during Viral Persistence. J. Virol., April 2017. ISSN 1098-5514. doi: 10.1128/JVI. 01083-16.

34. Emmanuelle Boulanger, Renan Duprez, Eric Delabesse, Jean Gabarre, Elizabeth Macintyre, and Antoine Gessain. Monololigoclonal pattern of Kaposi Sarcoma-associated herpesvirus (KSHV/HHV-8) episomes in primary effusion lymphoma cells. Int. J. Cancer, 115 (4):511–518, July 2005. ISSN 1097-0215. doi: 10.1002/ijc.20926.

35. Adam Grundhoff and Don Ganem. Inefficient establishment of KSHV latency suggests an additional role for continued lytic replication in Kaposi sarcoma pathogenesis. Journal of Clinical investigation, 113(1):124–136, January 2004. ISSN 0021-9738. doi: 10.1172/ JCI200417803.

36. Michael Stürzl, Dominika Gaus, Wilhelm G. Dirks, Don Ganem, and Ramona Jochmann. Kaposi’s sarcoma-derived cell line SLK is not of endothelial origin, but is a contaminant from a known renal carcinoma cell line. International Journal of Cancer, 132(8):1954–1958, April 2013. ISSN 00207136. doi: 10.1002/ijc.27849.

37. T. Piolot, M. Tramier, M. Coppey, J.-C. Nicolas, and V. Marechal. Close but Distinct Regions of Human Herpesvirus 8 Latency-Associated Nuclear Antigen 1 Are Responsible for Nuclear Targeting and Binding to Human Mitotic Chromosomes. Journal of Virology, 75(8):3948–3959, April 2001. ISSN 0022-538X. doi: 10.1128/JVI.75.8.3948-3959.2001.

38. John F. Domsic, Horng-Shen Chen, Fang Lu, Ronen Marmorstein, and Paul M. Lieberman. Molecular Basis for Oligomeric-DNA Binding and Episome Maintenance by KSHV LANA. PLoS Pathogens, 9(10):e1003672, October 2013. ISSN 1553-7374. doi: 10.1371/journal.ppat.1003672.

39. Alistair N. Boettiger, Bogdan Bintu, Jeffrey R. Moffitt, Siyuan Wang, Brian J. Beliveau, Geoffrey Fudenberg, Maxim Imakaev, Leonid A. Mirny, Chaoting Wu, and Xiaowei Zhuang. Super-resolution imaging reveals distinct chromatin folding for different epigenetic states. Nature, 529(7586):418–422, January 2016. ISSN 0028-0836, 1476-4687. doi: 10.1038/nature16496.

40. A. V. McDonnell, T. Jiang, A. E. Keating, and B. Berger. Paircoil2: improved prediction of coiled coils from sequence. Bioinformatics, 22(3):356–358, February 2006. ISSN 1367-4803. doi: 10.1093/bioinformatics/bti797.

41. Tuna Toptan, Lidia Fonseca, Hyun Jin Kwun, Yuan Chang, and Patrick S. Moore. Complex Alternative Cytoplasmic Protein Isoforms of the Kaposi’s Sarcoma-Associated Herpesvirus Latency-Associated Nuclear Antigen 1 Generated through Noncanonical Translation Initiation. J. Virol., 87(5):2744–2755, March 2013. ISSN 0022-538X, 1098-5514. doi: 10.1128/JVI.03061-12.

42. L.-Y. Wong and A. C. Wilson. Kaposi’s Sarcoma-Associated Herpesvirus Latency-Associated Nuclear Antigen Induces a Strong Bend on Binding to Terminal Repeat DNA. Journal of Virology, 79(21):13829–13836, October 2005. ISSN 0022-538X. doi: 10.1128/JVI.79.21.13829-13836.2005.

43. Rajesh Ponnusamy, Maxim V. Petoukhov, Bruno Correia, Tania F. Custodio, Franceline Juillard, Min Tan, Marta PiresdeMiranda, Maria A. Carrondo, J. Pedro Simas, Kenneth M. Kaye, Dmitri I. Svergun, and Colin E. McVey. KSHV but not MHV-68 LANA induces a strong bend upon binding to terminal repeat viral DNA. Nucl. Acids Res., 43(20):10039–10054, November 2015. ISSN 0305-1048, 1362-4962. doi: 10.1093/nar/gkv987.

44. Thomas Schalch, Sylwia Duda, David F. Sargent, and Timothy J. Richmond. X-ray structure of a tetranucleosome and its implications for the chromatin fibre. Nature, 436(7047):138–141, July 2005. ISSN 0028-0836, 1476-4679. doi: 10.1038/nature03686.

45. R Renne, W Zhong, B Herndier, M McGrath, N Abbey, D Kedes, and D Ganem. Lytic growth of Kaposi’s sarcoma-associated herpesvirus (human herpesvirus 8) in culture. Nature Medicine, 2(3):342–346, March 1996.

46. J. Menezes, W. Leibold, G. Klein, and G. Clements. Establishment and characterization of an Epstein-Barr virus (EBC)-negative lymphoblastoid B cell line (BJA-B) from an exceptional, EBV-genome-negative African Burkitt’s lymphoma. Biomedicine, 22(4):276–284, July 1975. ISSN 0300-0893.

47. K. F. Brulois, H. Chang, A. S.-Y. Lee, A. Ensser, L.-Y. Wong, Z. Toth, S. H. Lee, H.-R. Lee, J. Myoung, D. Ganem, T.-K. Oh, J. F. Kim, S.-J. Gao, and J. U. Jung. Construction and Manipulation of a New Kaposi’s Sarcoma-Associated Herpesvirus Bacterial Artificial Chromosome Clone. Journal of Virology, 86(18):9708–9720, September 2012. ISSN 0022-538X. doi: 10.1128/JVI.01019-12.

48. R. Renne, C. Barry, D. Dittmer, N. Compitello, P. O. Brown, and D. Ganem. Modulation of Cellular and Viral Gene Expression by the Latency-Associated Nuclear Antigen of Kaposi’s Sarcoma-Associated Herpesvirus. Journal of Virology, 75(1):458–468, January 2001. ISSN 0022-538X. doi: 10.1128/JVI.75.1.458-468.2001.

49. Paul Kellam, Dimitra Bourboulia, Nicolas Dupin, Chris Shotton, Cyril Fisher, Simon Talbot, Chris Boshoff, and Robin A. Weiss. Characterization of monoclonal antibodies raised against the latent nuclear antigen of human herpesvirus 8. Journal of virology, 73(6):5149–5155, 1999.

50. Samuel T. Hess, Thanu P.K. Girirajan, and Michael D. Mason. Ultra-High Resolution Imaging by Fluorescence Photoactivation Localization Microscopy. Biophysical Journal, 91(11): 4258–4272, December 2006. ISSN 00063495. doi: 10.1529/biophysj.106.091116.

51. Travis J Gould, Vladislav V Verkhusha, and Samuel T. Hess. Imaging biological structures with fluorescence photoactivation localization microscopy. Nature Protocols, 4(3):291–308, February 2009. ISSN 1754-2189,1750-2799. doi: 10.1038/nprot.2008.246.

52. Russell E Thompson, Daniel R Larson, and Watt W Webb. Precise nanometer localization analysis for individual fluorescent probes. Biophys J, 82(5):2775–2783, May 2002. ISSN 0006-3495.

53. B. Huang, W. Wang, M. Bates, and X. Zhuang. Three-Dimensional Super-Resolution Imaging by Stochastic Optical Reconstruction Microscopy. Science, 319(5864):810–813, February 2008. ISSN 0036-8075,1095-9203. doi: 10.1126/science.1153529.

54. S. Hell, G. Reiner, C. Cremer, and E. H. K. Stelzer. Aberrations in confocal fluorescence microscopy induced by mismatches in refractive index. Journal of Microscopy, 169(3):391–405, March 1993. ISSN 1365-2818. doi: 10.1111/j.1365-2818.1993.tb03315.x.

55. R Arimoto and J M Murray. A common aberration with water-immersion objective lenses. J Microsc, 216(Pt 1):49–51, October 2004. ISSN 0022-2720. doi: 10.1111/j.0022-2720.2004. 01383.x.

56. Stéfan van der Walt, S. Chris Colbert, and Gaël Varoquaux. The NumPy Array: A Structure for Efficient Numerical Computation. Computing in Science & Engineering, 13(2):22–30, March 2011. ISSN 1521-9615. doi: 10.1109/MCSE.2011.37.

57. Eric Jones, Travis Oliphant, Pearu Peterson, and others. SciPy: Open source scientific tools for Python, 2001.

58. Michael C. DeSantis, Shannon Kian Zareh, Xianglu Li, Robert E. Blankenship, and Y. M. Wang. Single-image axial localization precision analysis for individual fluorophores. Opt Express, 20(3):3057–3065, January 2012. ISSN 1094-4087. doi: 10.1364/ÜE.20.003057.

59. Manuel Guizar-Sicairos, Samuel T. Thurman, and James R. Fienup. Efficient subpixel image registration algorithms. Opt. Lett., OL, 33(2):156–158, January 2008. ISSN 1539-4794. doi: 10.1364/OL.33.000156.

60. Fabian Pedregosa, Gaël Varoquaux, Alexandre Gramfort, Vincent Michel, Bertrand Thirion, Olivier Grisel, Mathieu Blondel, Peter Prettenhofer, Ron Weiss, Vincent Dubourg, Jake Vanderplas, Alexandre Passos, David Cournapeau, Matthieu Brucher, Matthieu Perrot, and Édouard Duchesnay. Scikit-learn: Machine Learning in Python. Journal of Machine Learning Research, 12(Oct):2825–2830, 2011. ISSN ISSN 1533-7928.

61. Josef Perktold, Skipper Seabold, Jonathan Taylor, and statsmodels contributors. Statsmodels, 2017.

62. Philip J. Schneider and David H. Eberly. Geometric Tools for Computer Graphics. Morgan Kaufmann series in computer graphics and geometric modeling. Morgan Kaufmann, San Francisco, CA, USA, 2003. ISBN 1-55860-594-0978-1-55860-594-7.

63. John D. Hunter. Matplotlib: A 2d Graphics Environment. Computing in Science & Engineering, 9(3):90–95, May 2007. ISSN 1521-9615. doi: 10.1109/MCSE.2007.55.

64. P. Ramachandran and G. Varoquaux. Mayavi: 3d Visualization of Scientific Data. Computing in Science & Engineering, 13(2):40–51, 2011. ISSN 1521-9615.

65. Peter J. A. Cock, Tiago Antao, Jeffrey T. Chang, Brad A. Chapman, Cymon J. Cox, Andrew Dalke, Iddo Friedberg, Thomas Hamelryck, Frank Kauff, Bartek Wilczynski, De Hoon, and Michiel J. L. Biopython: freely available Python tools for computational molecular biology and bioinformatics. Bioinformatics, 25(11):1422–1423, June 2009. ISSN 1367-4803. doi: 10.1093/bioinformatics/btp163.

66. Thomas Hamelryck and Bernard Manderick. PDB file parser and structure class implemented in Python. Bioinformatics, 19(17):2308–2310, November 2003. ISSN 1367-4803. doi: 10.1093/bioinformatics/btg299.

67. Matthew Z. Tien, Dariya K. Sydykova, Austin G. Meyer, and Claus O. Wilke. PeptideBuilder: A simple Python library to generate model peptides. PeerJ, 1:e80, May 2013. ISSN 2167-8359. doi: 10.7717/peerj.80.

68. Schrödinger, LLC. The PyMOL Molecular Graphics System, Version 1.7, November 2015.

69. Christopher W. Wood, Marc Bruning, Amaurys Á Ibarra, Gail J. Bartlett, Andrew R. Thomson, Richard B. Sessions, R. Leo Brady, and Derek N. Woolfson. CCBuilder: an interactive web-based tool for building, designing and assessing coiled-coil protein assemblies. Bioinformatics, 30(21):3029–3035, November 2014. ISSN 1367-4803. doi: 10.1093/bioinformatics/btu502.

70. R. Byrd, P. Lu, J. Nocedal, and C. Zhu. A Limited Memory Algorithm for Bound Constrained Optimization. S/AMJ. Sc/. Comput., 16(5):1190–1208, September 1995. ISSN 1064-8275. doi: 10.1137/0916069.

